# Delivering large genes using adeno-associated virus and the CRE-lox DNA recombination system

**DOI:** 10.1101/2024.04.10.588864

**Authors:** Poppy Datta, Kun Do Rhee, Rylee J Staudt, Jacob M Thompson, Ying Hsu, Salma Hassan, Arlene V. Drack, Seongjin Seo

**Affiliations:** Department of Ophthalmology and Visual Sciences, The University of Iowa Carver College of Medicine, Iowa City, IA 52242, U.S.A; Institute for Vision Research, The University of Iowa Carver College of Medicine, Iowa City, IA 52242, U.S.A

## Abstract

Adeno-associated virus (AAV) is a safe and efficient gene delivery vehicle for gene therapies. However, its relatively small packaging capacity limits its use as a gene transfer vector. Here, we describe a strategy to deliver large genes that exceed the AAV’s packaging capacity using up to four AAV vectors and the CRE-lox DNA recombination system. We devised novel lox sites by combining non-compatible and reaction equilibrium-modifying lox site variants. These lox sites facilitate sequence-specific and near-unidirectional recombination of AAV vector genomes, enabling efficient reconstitution of up to 16 kb of therapeutic genes in a pre-determined configuration. Using this strategy, we have developed AAV gene therapy vectors to deliver *IFT140*, *PCDH15*, *CEP290*, and *CDH23* and demonstrate efficient production of full-length proteins in cultured mammalian cells and mouse retinas. Notably, this approach significantly surpasses the trans-splicing and split-intein-based reconstitution methods in efficiency, requiring lower doses, minimizing or eliminating the production of truncated protein products, and offering flexibility in selecting splitting positions. The CRE-lox approach described here provides a simple and effective platform for producing AAV gene therapy vectors beyond AAV’s packaging capacity.

## Introduction

Adeno-associated virus (AAV) is one of the most broadly used viral vectors for gene therapy. With diverse natural and engineered serotypes, AAVs can transduce a wide range of cell types, while eliciting minimal immune responses (1–4). Furthermore, AAV-mediated gene delivery appears to result in long-term transgene expression, which is an important qualification for a cure for a genetic disease. The safety, broad tropism, and long-term transgene expression have made AAV the preferred vector for gene therapy in numerous clinical trials.

However, AAV has a major disadvantage as a gene transfer vector. AAVs can accommodate up to 5 kb of viral genomes (5), and this packaging capacity is insufficient to carry full-length coding sequences of many disease-causing genes. For instance, the full-length coding sequence of *ABCA4*, which is the most frequently mutated gene among all Mendelian retinal degeneration genes (6), spans 6,822 bp. A ciliopathy gene *CEP290* is associated with multiple genetic diseases ranging from isolated early-onset retinal degeneration to oculorenal dysplasia and lethal Meckel-Gruber syndrome (7–11). Mutations in *CEP290* are the most frequent cause of Leber congenital amaurosis (LCA), severe vision loss at birth or in infancy, and its coding sequence is 7,440 bp-long. The coding sequence of *CDH23*, mutations of which cause Usher syndrome affecting both vision and hearing, spans 10,065 bp (12, 13). Since viral packaging signals (i.e., inverted terminal repeats (ITRs)) and other essential regulatory elements (e.g., a promoter and a transcription termination signal) must be included in viral vectors, the practical limit of a therapeutic gene that may be delivered via AAVs is ∼4 kb. This limited packaging capacity of AAV precludes its use as a gene transfer vector when large gene delivery is needed.

Various strategies have been developed to deliver genes larger than the packaging capacity of AAV vectors (14, 15). In these strategies, cargo genes are split and packaged in two (or three) AAV vectors for delivery to target cells. The split genes can be reconstituted at the DNA, mRNA, or protein level. The DNA-level reconstitution approach relies on occasional DNA recombination events that occur between AAV genomes after internalization. This strategy includes dual (or triple) AAV trans-splicing and hybrid methods (16–19). A new method to combine two AAV vector genomes using the CRE-lox DNA recombination system has been recently reported (20). In these approaches, the reconstituted therapeutic cassette produces mRNAs encoding full-length proteins. Additionally, a novel strategy, REVeRT (<u>re</u>constitution <u>v</u>ia m<u>R</u>NA trans-splicing), has been reported to facilitate the reconstitution of large genes at the mRNA level (21). The protein-level reconstitution approach employs split inteins and protein trans-splicing (22–24). Protein splicing is an auto-catalytic process where an intervening protein segment (intein) excises itself from a precursor protein and attaches its two flanking regions (exteins) through a peptide bond. Some inteins are naturally split, originating from two separate genes (23, 24), and the protein trans-splicing approach utilizes these naturally split inteins. In this approach, the N-and C-terminal halves of a protein are separately delivered and produced from corresponding AAV vectors, and reconstitution is achieved via protein trans-splicing mediated by split inteins.

Although these approaches have produced encouraging outcomes in animal models (17, 19, 21, 25–29), the need for more efficient and versatile delivery methods remains. For instance, AAV trans-splicing methods require high-dose AAV injections to compensate for their low efficiency and the random configuration of recombination events. This problem becomes more pronounced when more than two AAV vectors are needed. Similarly, the reconstitution efficiency of REVeRT appears to decrease significantly when the cargo gene is split into three AAV vectors. The split intein-based reconstitution approach entails the production of truncated proteins, which could potentially have dominant negative effects. Furthermore, the efficiency of protein splicing is heavily affected by the amino acids immediately adjacent to split inteins (24, 30, 31), and the impact of truncation on the folding and stability of both truncated protein products and reconstituted proteins is hard to predict. Due to these constraints, determining optimal splitting positions requires a screening process, and suitable positions may not always be available.

In the present work, we describe a novel strategy to deliver large genes using up to four AAV vectors. Cargo genes are split into 2-4 AAV vectors and reconstituted by using the CRE-lox DNA recombination system. The use of novel lox sites, which were generated by combining non-compatible and reaction equilibrium-modifying lox site variants, enables efficient reconstitution of a therapeutic cassette in a pre-determined configuration.

## Results

### Development of novel lox site variants for sequence-specific, unidirectional recombination

The canonical loxP site consists of two 13-bp inverted repeats (left and right elements; LE and RE, respectively) separated by an asymmetric 8-bp spacer/core sequence (**Figure 1A**). While the left and right elements serve as the binding sites for the CRE recombinase, the spacer participates in the strand exchange reaction and determines the compatibility between lox sites (i.e., whether two lox sites can recombine or not) (32–34). The asymmetry of the spacer imparts directionality to the loxP site.

**Figure 1.**
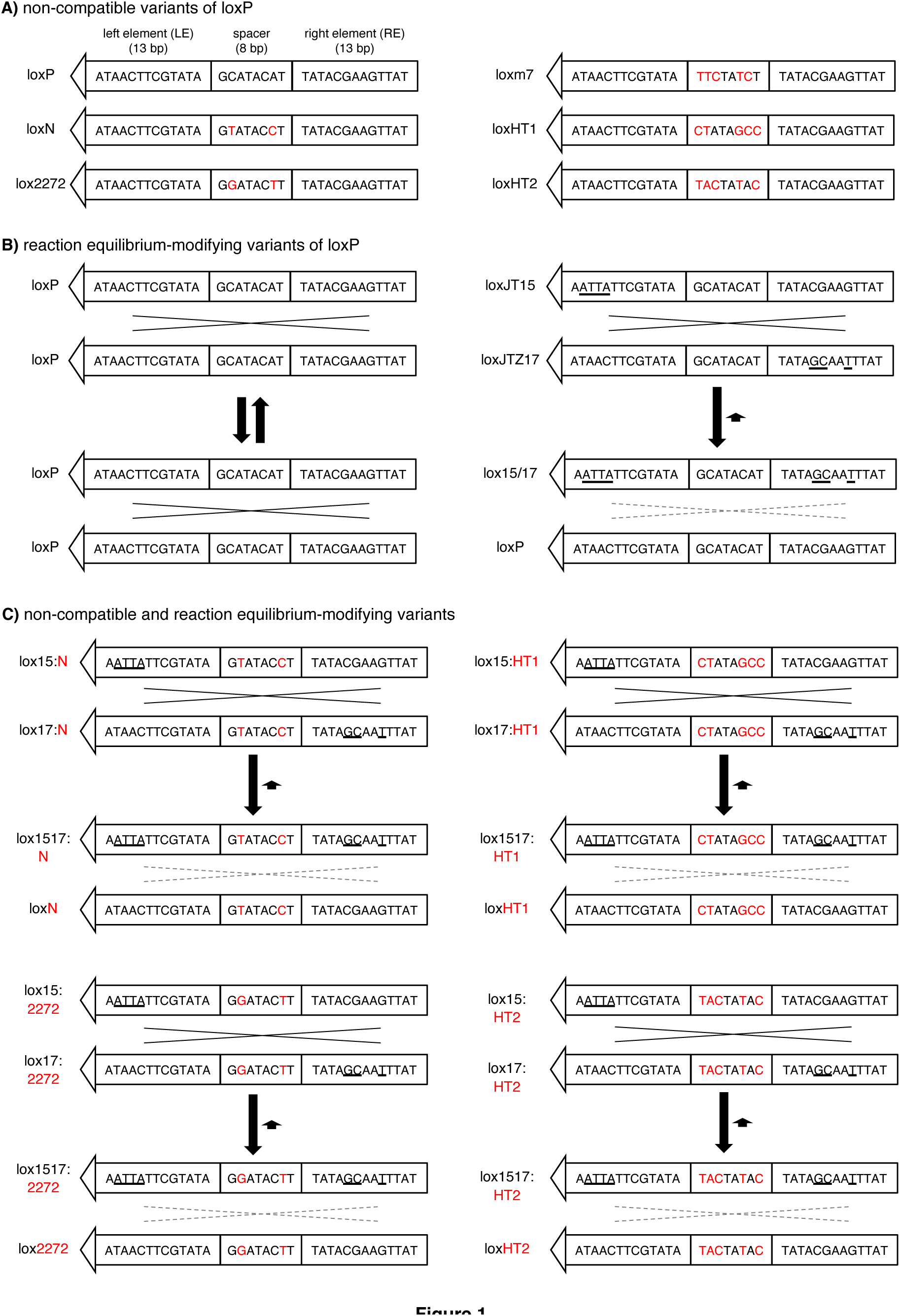
Lox site variants that enable CRE-dependent splicing of multiple AAV vector genomes. (A) Non-compatible mutant variants of loxP. Sequence differences (red) in the spacer region prevent recombination between non-compatible lox sites. Left and right elements (LE and RE, respectively) are palindromic. (B) Reaction equilibrium-modifying variants of loxP. LoxJT15 and loxJTZ17 sites have mutations (underlined) in either LE or RE but not in both. Recombination between two loxP sites (left) is fully reversible because substrates and products are identical. In contrast, recombination between loxJT15 and loxJTZ17 produces an LE/RE double mutant (lox15/17) and a loxP sequence. LE/RE double mutants are poor substrates for CRE, and consequently, the reverse reaction is significantly slower than the forward reaction. (C) Lox site variants that prevent recombination between non-compatible lox sites and inhibit reverse reactions. The spacer sequences of loxJT15 and loxJTZ17, which are from loxP, are replaced with those of the non-compatible lox sites (loxN, lox2272, loxm7 (not shown), loxHT1, and loxHT2).

There are two classes of lox site variants. One is non-compatible mutant variants, which include loxm7, loxN, and lox2272 (35–37) (**Figure 1A**). These variants have mutations within the spacer sequence, and these mutations prevent strand exchange (and consequently recombination) between non-compatible lox sites while allowing recombination between homologous (or compatible) sites. A high-throughput screen identified fully non-compatible and promiscuous lox sites (38). The second group comprises reaction equilibrium-modifying variants (**Figure 1B**). These variants have mutations in either LE or RE, but not in both (e.g., loxJT15, loxJTZ17, lox71, and lox66) (39, 40). Single-element mutations do not affect the binding of CRE to the lox site, and recombination between these mutant lox sites occurs as efficiently as between canonical loxP sites. However, recombination between LE and RE single mutants produces an LE/RE double mutant and a canonical loxP site. The presence of mutations in both LE and RE significantly reduces the affinity of the LE/RE double mutant to CRE, making it a poor substrate. While the recombination between canonical loxP sites is fully reversible as the initial substrates and the products have the same lox sites, the reaction equilibrium is drastically shifted toward the forward direction when LE and RE single mutants are used as substrates because the reverse reaction is much slower than the forward reaction. This causes the CRE-mediated recombination nearly unidirectional when the reaction equilibrium-modifying lox sites are used.

Although the CRE-lox system is highly efficient, canonical loxP sites (or any single species of lox sites) cannot be used to combine more than two DNA molecules. When there are two or more loxP sites in a single DNA molecule, the intervening “floxed” sequence is rapidly excised (**Figure S1A**). The reverse reaction (i.e., insertion) is much slower than the forward reaction. Furthermore, if all DNA fragments have the same lox sites, recombination can occur in any combination (**Figure S1B**), preventing the specification of the order of DNA segments in end products, leading to the production of unintended, non-functional products. Lastly, since the CRE-lox recombination is fully reversible, the reconstituted DNA constantly goes through the assembly-disassembly cycle, limiting the yield of the reconstituted DNAs (**Figure S1C**). In principle, the yields of reconstituted products are 50%, 25%, and 12.5% when 2, 3, and 4 fragments are used as substrates, respectively, even when the excision and the order of DNA segment problems are disregarded.

To enable the reconstitution of large genes using multiple AAV vectors and the CRE-lox DNA recombination system, we devised novel lox sites by combining non-compatible and reaction equilibrium-modifying lox site variants (**Figure 1C**). We chose the loxJT15-loxJTZ17 pair for the reaction equilibrium-modifying mutants because this pair was the most effective in inhibiting reverse-direction recombination (39). For the non-compatible lox sites, we selected loxN, lox2272, loxm7, and two additional lox sites identified by a high-throughput screen with spacer sequences CTATAGCC (named loxHT1 herein) and TACTATAC (loxHT2) (38). The loxN-based pair, for example, was generated by replacing the spacer sequence of loxJT15 and loxJTZ17 (GCATACAT) with that of loxN (GTATACCT). If these hybrid lox sites are fully non-compatible with one another, they should prevent the excision of intervening sequences and unintended recombinations (**Figure S1D and E**). At the same time, they should significantly increase the yield of reconstituted DNA products by inhibiting reverse reactions, particularly when using 3 or more AAV vectors (**Figure S1F**).

To assess the compatibility of the developed hybrid lox sites in mammalian cells, we designed GFP expression cassettes capable of tracking recombination events between different lox sites (**Figure 2** and **Figure S2**; see **Supplementary Materials** for sequences). The first reporter construct, loxP-2272, was designed to survey the compatibility of loxJT15 with four hybrid lox sites (**Figure 2A**). The reporter is composed of a CMV promoter, a GFP coding sequence, a 156-bp segment from the human CEP290 C-terminus (C290C; amino acids 2428-2479; in frame with GFP), and a stop codon. A loxJT15 site (15:P) was inserted between GFP and C290C such that recombination events between loxJT15 and any downstream lox sites would result in the excision of C290C and the production of new GFP fusion proteins with an identification tag (**Figure S2**). FLAG, HA, MYC, and V5 tags were used to report the recombination of loxJT15 with loxJTZ17:m7 hybrid (hereafter denoted as 17:m7 for brevity), 17:HT1, 17:HT2, and 17:2272 sites, respectively. Recombination between loxJT15 (15:P) and 17:HT1, for instance, results in the production of GFP+HA fusion proteins (∼30 kDa). Of note, recombination events not involving the 15:P site, such as between 17:m7 and 17:2272, do not lead to the fusion of associated tags with GFP and therefore go unreported. Four additional reporter constructs (loxP-N, lox2272-N, loxP-HT2, and lox2272-HT2) were generated to examine the compatibility among the hybrid lox sites that we have developed (**Figure 2A**).

**Figure 2.**
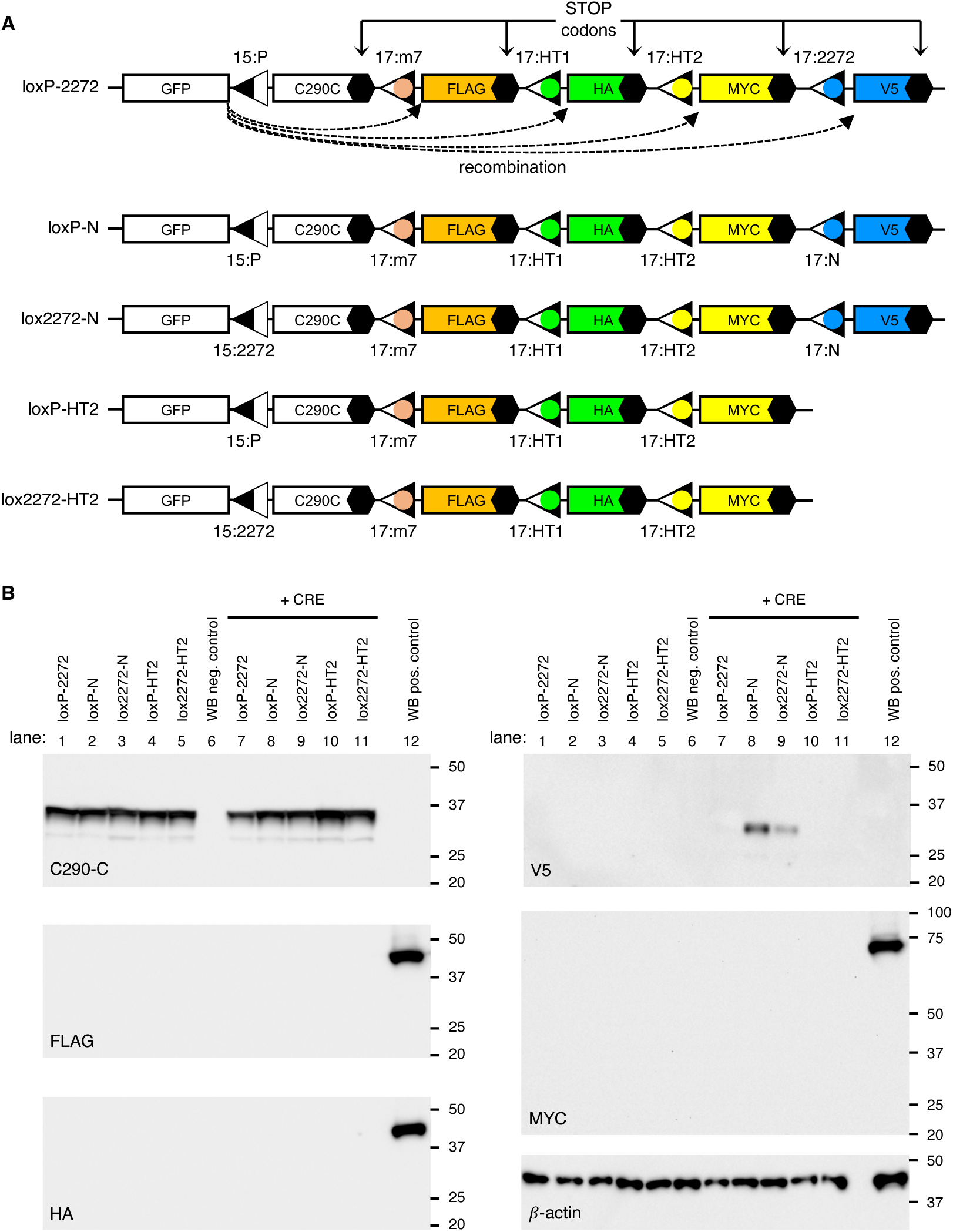
Assessment of incompatibility among hybrid lox sites. (A) Schematics of reporter constructs to detect recombination events between loxJT15 (15:P), loxJTZ17:m7 (17:m7), loxJTZ17:HT1 (17:HT1), loxJTZ17:HT2 (17:HT2), loxJTZ17:2272 (17:2272), and loxJTZ17:N (17:N). The names of the reporter constructs (loxP-2272, loxP-N, lox2272-N, loxP-HT2, and lox2272-HT2) are shown on the left. Black hexagons denote stop codons. C290C: a 156-bp fragment from human CEP290 C-terminus (aa 2428-2479). (B) The spacers of loxP and lox2272 are fully incompatible with each other and with those of loxm7, loxHT1, and loxHT2. Reporter constructs shown in (A) were transfected to HEK293T cells with and without a CRE expression vector, and cell lysates were subjected to SDS-PAGE and immunoblotting. C290-C, FLAG, HA, V5, MYC, and ý-actin antibodies were used for immunoblotting. A lysate derived from untransfected cells served as the negative control (lane 6), while lysates obtained from cells transfected with MYC-BBS1, FLAG-LZTFL1, and HA-LZTFL1 expression vectors were used as the positive control (lane 12). ý-actin was used as a loading control.

When transfected into HEK293T cells, these reporters produced 35-kDa GFP+C290C fusion proteins, which could be detected by our CEP290 antibody (**Figure 2B** and **Figure S2**). However, when co-transfected with a *CRE* expression plasmid (pAAV-EF1α-*CRE*), reporters containing the 17:N site (i.e., loxP-N and lox2272-N) expressed GFP+V5 fusion proteins. In contrast, no new GFP fusion proteins were detected in cells transfected with loxP-2272, loxP-HT2, and lox2272-HT2. These results suggest that the spacer of loxN is partially compatible with those of loxP and lox2272, while loxP and lox2272 are fully incompatible with each other and with loxm7, loxHT1, and loxHT2. Since these reporters are designed to disclose only recombination events that involve the first lox site, which is linked to GFP, the compatibility among downstream lox sites (i.e., loxm7, loxHT1, and loxHT2) was not tested in this assay. As a positive control for Western blotting (lane 12), lysates from cells transfected with FLAG-LZTFL1, HA-LZTFL1, and MYC-BBS1 expression vectors were included to rule out the possibility of Western blotting failure, with β-actin serving as a loading control. Based on these results, we selected the spacer sequences of loxP, lox2272, and loxHT1 for the assembly of up to four AAV vector genomes. The spacers of loxm7 and loxHT2 may be used as alternatives to loxHT1 in this assembly strategy.

### CRE-lox-mediated reconstitution of large genes delivered by tripartite AAV vectors

To evaluate the feasibility of our approach in delivering large genes using tripartite AAV vectors, we designed a set of three AAV vectors containing the coding sequences of three human genes: 5’ 1,923 bp of *IFT140* (*IFT140-N*), full-length *BBS1*, and full-length *LZTFL1* (**Figure 3A**). The first vector comprises a CMV promoter, 5’ 1,923 bp of the *IFT140* coding sequence, a splice donor (SD) site, and a loxJT15 site. An HA tag sequence was added to the 5’ end of *IFT140* to facilitate the detection of expressed proteins. The second vector contains a loxJTZ17 site, a splice acceptor (SA) site, the coding sequence of *BBS1* (1,775 bp; including an HA tag and two linker sequences), an SD site, and a lox15:2272 site. The third vector is composed of a lox17:2272 site, a SA site, the *LZTFL1* coding sequence (988 bp; including a linker sequence), and a bovine growth hormone (BGH) transcription termination signal. Recombination between these three AAV vector genomes in the correct arrangement will lead to the reconstitution of an expression cassette encoding IFT140N+BBS1+LZTFL1 fusion proteins.

**Figure 3.**
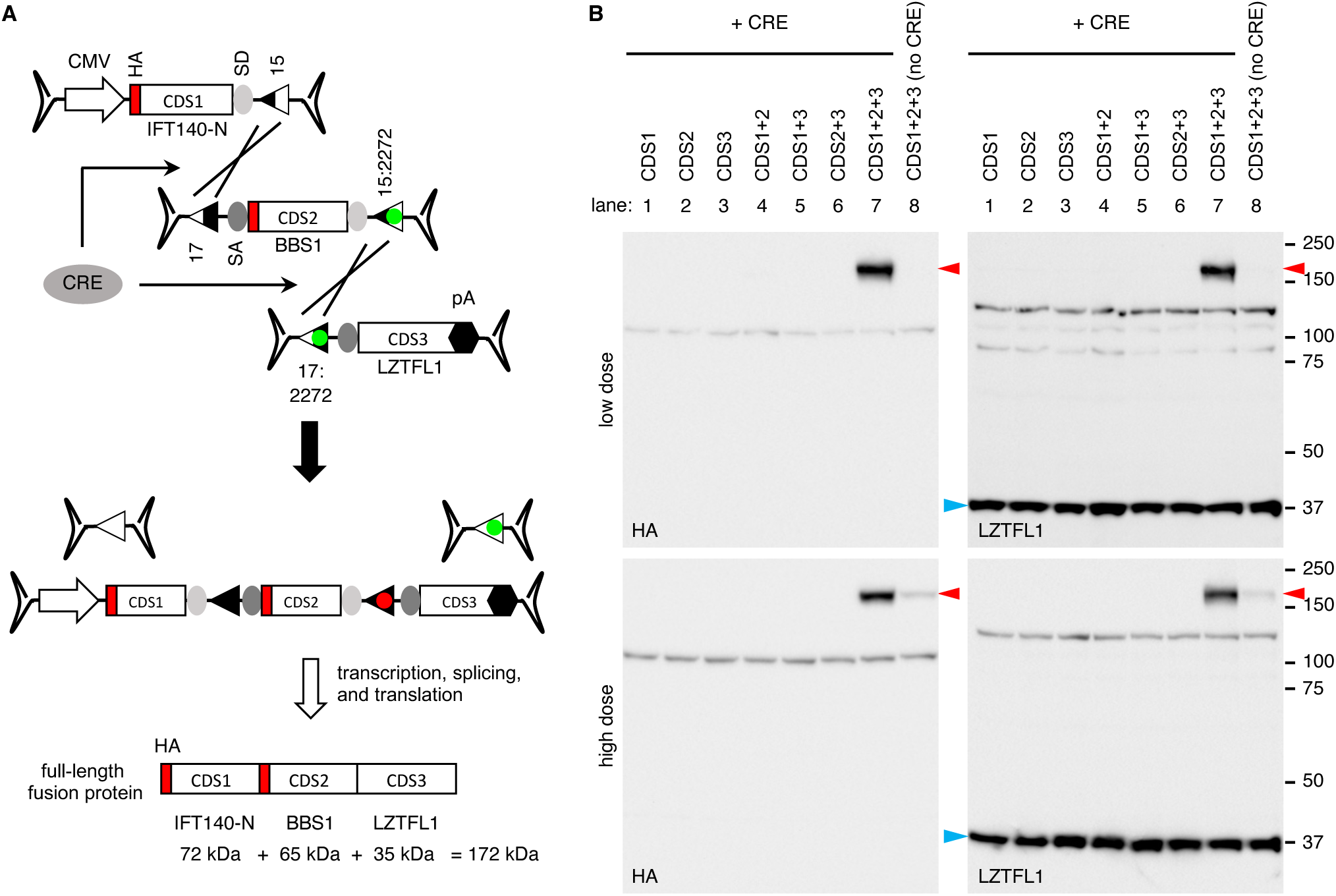
CRE-lox-mediated reconstitution of large genes delivered by tripartite AAV vectors: proof-of-concept. (A) Schematic representation of large gene reconstitution by CRE-lox mediated recombination of three AAV vector genomes. A gene-of-interest is split into three fragments (CDS1, 2, and 3) and delivered to target cells via three separate AAV vectors. For a proof-of-concept demonstration, *IFT140-N* (5’ 1,923 bp of *IFT140*), *BBS1,* and *LZTFL1* coding sequences were used as CDS1, 2, and 3, respectively. HA tag sequences were added to the 5’ ends of *IFT140-N* and *BBS1*. CRE recombinase was delivered via a separate AAV vector. SD: splice donor site, SA: splice acceptor site, 15: loxJT15, 17: loxJTZ17, and pA: polyA signal. (B) Production of IFT140N+BBS1+LZTFL1 fusion proteins (red arrowheads) from a tripartite AAV vector set. AAV vectors depicted in panel (A) were transduced to 293T cells, and the expression of IFT140N+BBS1+LZTFL1 fusion proteins was examined by SDS-PAGE and immunoblotting using HA and LZTFL1 antibodies. Numbers on the right mark the location of protein standards. A separate AAV vector, AAV-EF1α-CRE, was co-transduced to express CRE (lanes 1-7). Lane 1: CDS1 (IFT140-N) only, lane 2: CDS2 (BBS1) only, lane 3: CDS3 (LZTFL1) only, lane 4: CDS1 + CDS2, lane 5: CDS1 + CDS3, lane 6: CDS2 + CDS3, lane 7: CDS1 + CDS2 + CDS3, lane 8: CDS1 + CDS2 + CDS3 (without CRE). Endogenous LZTFL1 (blue arrowheads) was used as a loading control.

These AAV vectors, all utilizing serotype 2, were transduced individually or in various combinations to HEK293T cells at two different doses. The “low” dose involved transduction at a multiplicity of infection (MOI) of 1.5×10^4^ for each vector, while the “high” dose utilized an MOI of 6.0×10^4^ for each vector. CRE recombinase was delivered via a separate AAV vector (AAV2/2-EF1α-CRE) with an MOI of 0.3×10^4^ for the low dose and 1.2×10^4^ for the high dose. As shown in **Figure 3B**, robust expression of the IFT140N+BBS1+LZTFL1 fusion protein (red arrowheads) was observed using both HA and LZTFL1 antibodies at both low and high doses (upper and lower panels, respectively). The migration rate of the protein was consistent with the predicted molecular weight of the full-length fusion protein, approximately 172 kDa. Notably, no protein production was detected when the three AAV vectors were transduced individually or any of the three was omitted. Although the expression of IFT140N+BBS1+LZTFL1 fusion proteins was not detected in the absence of CRE (lane 8) at the low dose, a small amount was detectable at the high dose. These proteins are likely attributed to spontaneous, random recombination of AAV vector genomes occurring after internalization. Endogenous LZTFL1, marked by blue arrowheads, served as a loading control. These data indicate that the CRE-and hybrid lox site-mediated reconstitution is significantly more efficient than the trans-splicing approach for reconstituting large genes.

### CRE-lox-mediated reconstitution of large genes delivered by quadripartite AAV vectors

We expanded our approach to utilize quadripartite AAV vectors. For a proof-of-concept demonstration, we designed AAV vectors containing the coding sequences of the 5’ 1,923 bp of *IFT140*, *IFT57*, *BBS5*, and *LZTFL1* (**Figure 4A**). The first and fourth AAV vectors that contained the *IFT140-N* and *LZTFL1* coding sequences were the same ones used in the tripartite set above. The second vector was constructed with a loxJTZ17 site, a SA site, the *IFT57* coding sequence (1,330 bp; including two linker sequences), a SD site, and a lox15:HT1 site. The third vector was composed of a lox17:HT1 site, a SA site, the *BBS5* coding sequence (1,090 bp; including an HA-tag and two linker sequences), a SD site, and a lox15:2272 site. If recombination occurs as intended, it will result in the reconstitution of an expression cassette encoding IFT140N+IFT57+BBS5+LZTFL1 fusion proteins with a predicted molecular weight of ∼200 kDa.

**Figure 4.**
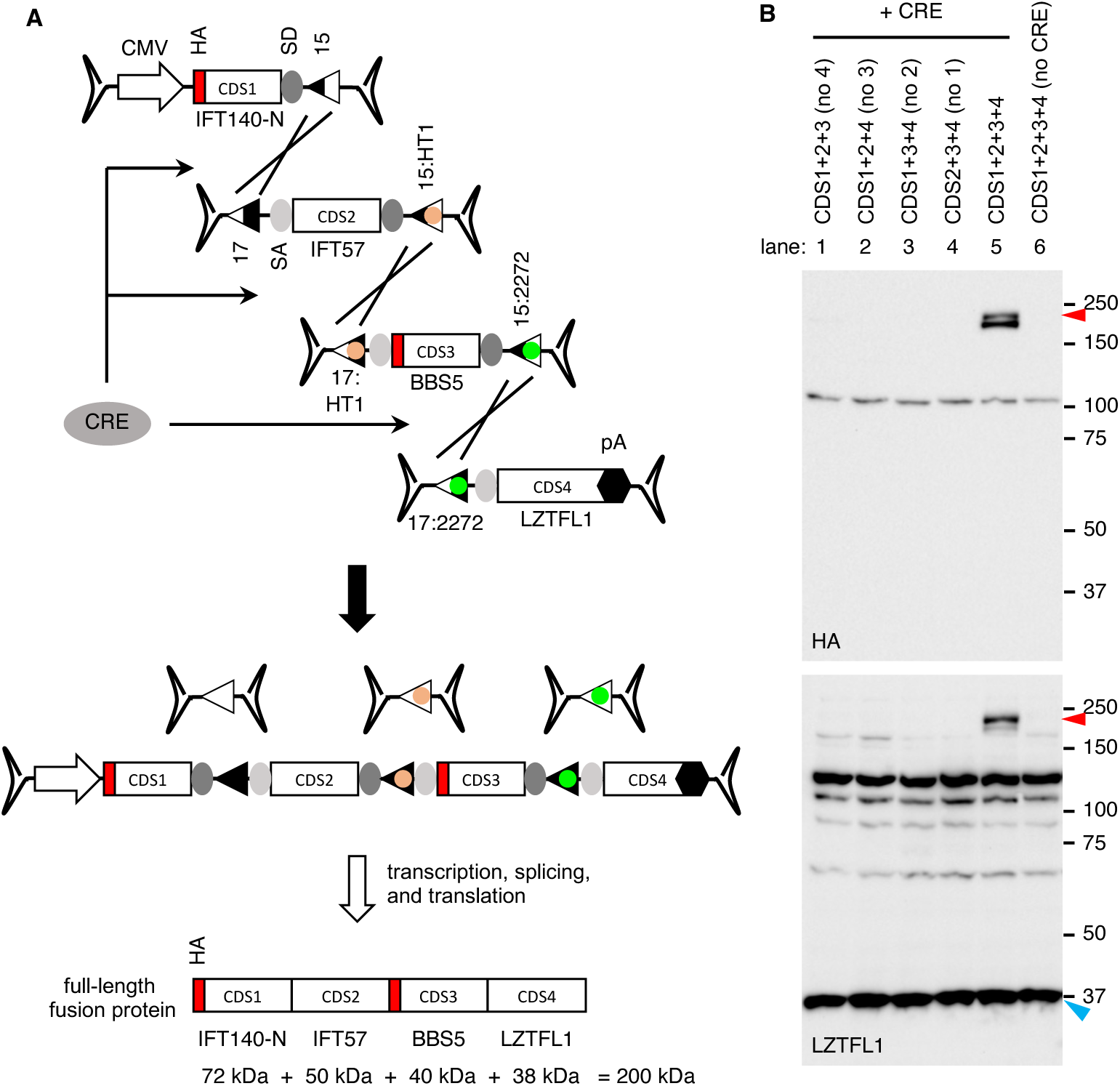
CRE-lox-mediated reconstitution of large genes delivered by quadripartite AAV vectors: proof-of-concept. (A) Schematic representation of large gene reconstitution by CRE-lox mediated recombination of four AAV vector genomes. A gene-of-interest is split into four fragments (CDS1, 2, 3, and 4) and delivered to target cells via four separate AAV vectors. For a proof-of-concept demonstration, *IFT140-N* (5’ 1,923 bp of *IFT140*), *IFT57, BBS5,* and *LZTFL1* coding sequences were used as CDS1, 2, 3, and 4, respectively. HA tag sequences were added to the 5’ ends of *IFT140-N* and *BBS5*. Others are the same as in Figure 3. (B) Production of IFT140N+IFT57+BBS5+LZTFL1 fusion proteins (red arrowheads) from a quadripartite AAV vector set. AAV vectors depicted in panel (A) were transduced to 293T cells, and the expression of IFT140N+IFT57+BBS5+LZTFL1 fusion proteins was examined by SDS-PAGE and immunoblotting using HA and LZTFL1 antibodies. Endogenous LZTFL1 (blue arrowheads) was used as a loading control. Others are the same as in Figure 3.

These AAV vectors were transduced into 293T cells in various combinations at an MOI of 2.5×10^4^ per vector. The AAV-EF1α-CRE vector was transduced at an MOI of 0.5×10^4^. Upon transduction of the quadripartite AAV vectors and AAV-EF1α-CRE, the production of IFT140N+IFT57+BBS5+LZTFL1 fusion proteins was readily detected by immunoblotting using HA and LZTFL1 antibodies (**Figure 4B**; lane 5). The fusion protein appeared to be unstable, with some instances of truncation near the C-terminus, resulting in doublets. When any of the four AAV vectors was omitted, the full-length fusion protein was not produced (lanes 1-4). Furthermore, in the absence of CRE, the fusion protein was not detected (lane 6). These data validate the successful reconstitution of a gene expression cassette delivered by quadripartite AAV vectors and underscore the efficacy of our CRE-lox-mediated recombination approach.

### CRE-lox-mediated reconstitution of IFT140 delivered by bipartite AAV vectors

We applied the CRE-lox-mediated DNA recombination approach to *IFT140*, a gene associated with retinitis pigmentosa (RP) and short-rib thoracic dysplasia (41–43). Although the full-length *IFT140* coding sequence (4,389 bp) is small enough to be accommodated within a single AAV vector, additional regulatory sequences such as a promoter, a transcription termination signal, and two inverted terminal repeats (ITRs) must be included in the gene therapy vector, and the addition of such sequences makes the *IFT140* expression cassette to exceed the AAV’s packaging capacity. Therefore, at least two AAV vectors are required to deliver the *IFT140* gene. Among various dual AAV approaches developed to date, the split intein-mediated protein trans-splicing method has demonstrated notable effectiveness (28, 29, 44). Among split inteins examined, gp41 split inteins are one of the most efficient in facilitating protein trans-splicing ((28, 45, 46) and our unpublished data). We constructed dual AAV-*IFT140* vector sets with the CRE-lox-mediated DNA recombination approach and the gp41 split intein-mediated protein trans-splicing method and compared their efficiencies in producing full-length proteins.

The first set, designed for the CRE-lox approach (**Figure 5A**), consists of the 5’ vector that contains a CMV (or CBh) promoter, 5’ 1,923 bp of the *IFT140* coding sequence, an SD site, and a lox15:2272 site and the 3’ vector that contains a lox17:2272 site, an SA site, the rest of the *IFT140* coding sequence (2,466 bp), and a BGH transcription termination signal. Since the combined payload capacity of two AAV vectors is ∼9 kb and the size of the *IFT140* expression cassette is 6-6.5 kb, there is space to accommodate the *CRE* gene within the dual AAV-*IFT140* vectors. We incorporated the *CRE* coding sequence, along with an N-terminal T2A “self-cleaving” peptide, after the lox15:2272 site in the 5’ vector. A BGH polyA signal was also added following the *CRE* gene. The inclusion of *CRE* in this configuration not only eliminates the need for a separate AAV vector for *CRE* delivery but also offers an additional benefit of CRE “self-inactivation” by causing the separation of the *CRE* CDS from its promoter as recombination progresses.

**Figure 5.**
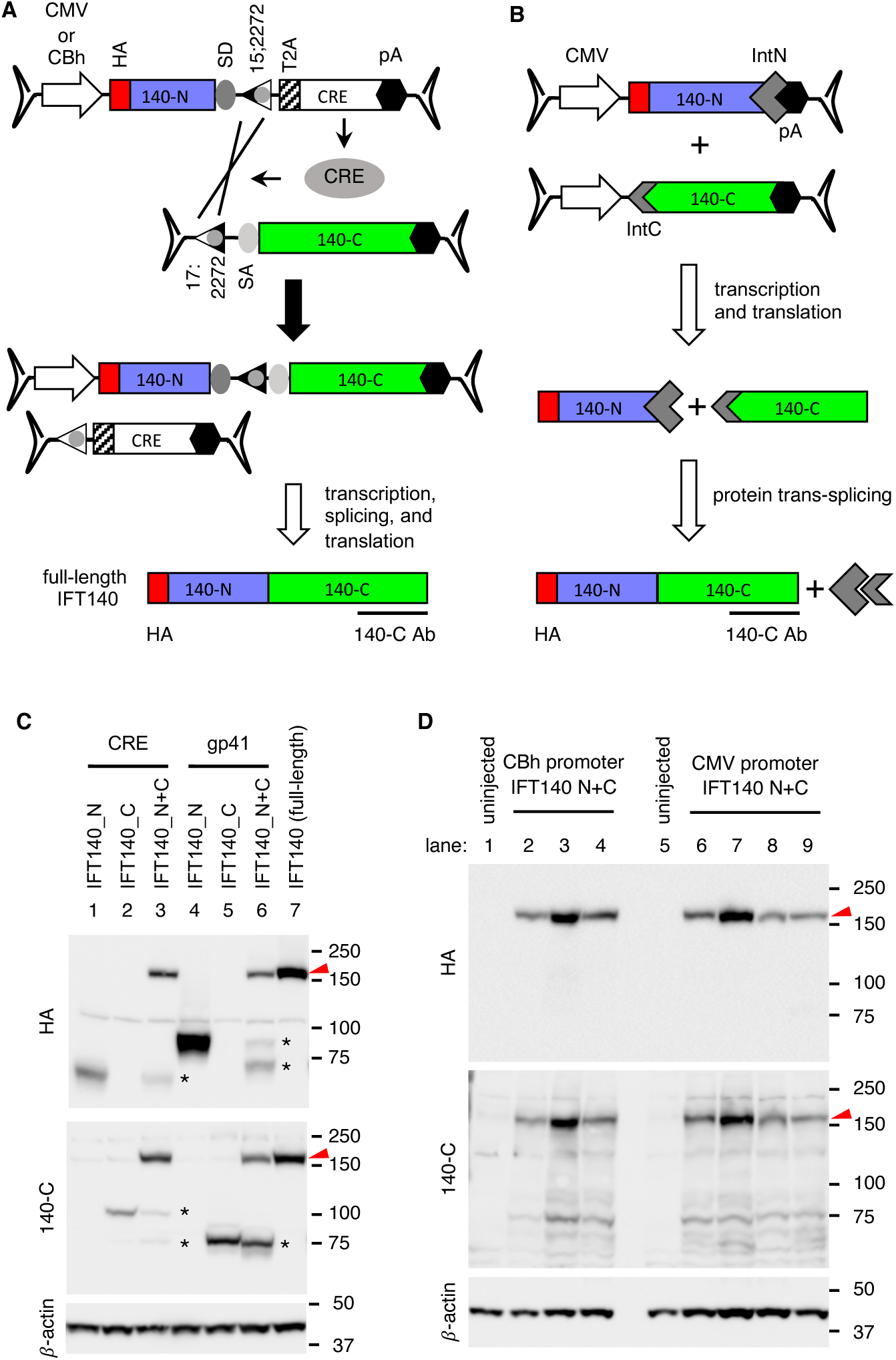
Reconstitution of IFT140 by CRE-lox-mediated recombination and protein trans-splicing. (A) Schematics of IFT140 reconstitution by CRE-lox-mediated recombination of bipartite AAV vector genomes. T2A: T2A “self-cleaving” peptide. Others are the same as in Figure 3. (B) Schematics of IFT140 reconstitution by gp41 split intein-mediated protein trans-splicing. The immunogen part used to raise the 140-C antibody was marked by a solid line at the bottom. IntN: N-terminal gp41 split intein, IntC: C-terminal gp41 split intein. (C) Production of full-length IFT140 proteins (red arrowheads) by CRE-lox-mediated recombination and protein trans-splicing approaches in 293T cells. HEK293T cells were transduced with dual AAV vectors depicted in panels A and B (with a CMV promoter), and cell lysates were subjected to SDS-PAGE followed by immunoblotting with HA and 140-C antibodies. Asterisks indicate truncated IFT140 protein products. Lanes 1-3: dual AAV-IFT140 CRE-lox set (with a CMV promoter), lane 1: AAV-IFT140_N only, lane 2: AAV-IFT140_C only, lane 3: dual AAV-IFT140_N+C, lanes 4-6: dual AAV-IFT140 gp41 set, lane 4: AAV-IFT140N-IntN only, lane 5: AAV-IFT140C-IntC only, lane 6: dual AAV-IFT140_N+C gp41, and lane 7: pCS2HA-IFT140 plasmid transfected (full-length; positive control). (D) Expression of full-length IFT140 from dual AAV-*IFT140* CRE/lox sets in mouse retinas. Dual AAV-*IFT140* vectors illustrated in panel A were administered via subretinal injection into mouse eyes (serotype: AAV5, dose: 5×10^9^ GC per vector), and retinal protein extracts were subjected to immunoblotting analysis. AAV-IFT140_N vectors with both CBh and CMV promoters were injected to explore potential differences in expression levels. Lysates from uninjected eyes were used as a negative control (lanes 1 and 5). ý-actin was used as a loading control.

For the protein trans-splicing approach, we chose to split the IFT140 protein at amino acids D767/C768 (**Figure S3**). This splitting position was determined based on the IFT140 domain structure and the amino acid residues known to support protein splicing (31). The 5’ vector of the split intein set (**Figure 5B**) consists of a CMV promoter, 5’ 2,301 bp of the *IFT140* coding sequence, the N-terminal gp41 split intein (IntN), and a BGH transcription termination signal. The 3’ vector of this set contains the same CMV promoter, the C-terminal gp41 split intein (IntC), the rest of the *IFT140* coding sequence (2,088 bp), and a BGH transcription termination signal. To facilitate protein detection, an HA tag was introduced at the N-terminus of IFT140 in both sets. Additionally, we used an IFT140 antibody (140-C Ab), which was raised against human IFT140 aa1114-1462, to detect the C-terminal portion of the protein.

Both sets of dual AAV-*IFT140* vectors (serotype 2) were transduced into 293T cells at an MOI of 3×10^4^ for each vector. Upon transduction, both approaches demonstrated efficient production of full-length IFT140 proteins (**Figure 5C**; lanes 3 and 6; red arrowheads). However, the split intein set exhibited significantly more truncated protein products compared to the CRE-lox set (lanes 3 and 6; asterisks). These results indicate that, when an equal amount of AAV vectors is transduced, the CRE-lox-mediated recombination approach is at least as effective as the protein trans-splicing approach in producing full-length proteins. Furthermore, the CRE-lox method has the additional advantage of generating little to no truncated protein products. As a positive control, a plasmid DNA encoding full-length IFT140 with an N-terminal HA tag was transfected and included (lane 7).

We further investigated whether the CRE/lox-based dual AAV-*IFT140* vectors could produce full-length IFT140 proteins in mouse retinas. To this end, wild-type mouse eyes underwent subretinal administration with two sets of CRE/lox dual AAV-*IFT140* vectors: one with a CMV promoter and the other with a CBh promoter. Both sets of AAV vectors were prepared with the AAV5 serotype and administered at a dose of 5×10^9^ genome copies (GC) per vector. Treated and some of the contralateral control eyes were collected three weeks post-injection and the production of full-length IFT140 proteins was examined by immunoblotting. As shown in **Figure 5D**, robust expression of full-length IFT140 was detected in all animals injected. These data highlight the reliability of the CRE/lox-mediated DNA reconstitution approach for delivering large genes using AAV.

### Reconstitution of PCDH15 by CRE-lox-mediated recombination

Mutations in *PCDH15* cause Usher syndrome type 1F (USH1F), which is characterized by profound congenital hearing impairment and progressive vision loss (47–49). The full-length human *PCDH15* CDS spans 5,865 bp, necessitating two AAV vectors for delivery. We applied the CRE-lox and split intein approaches to *PCDH15* and created dual AAV vectors (**Figure 6**).

**Figure 6.**
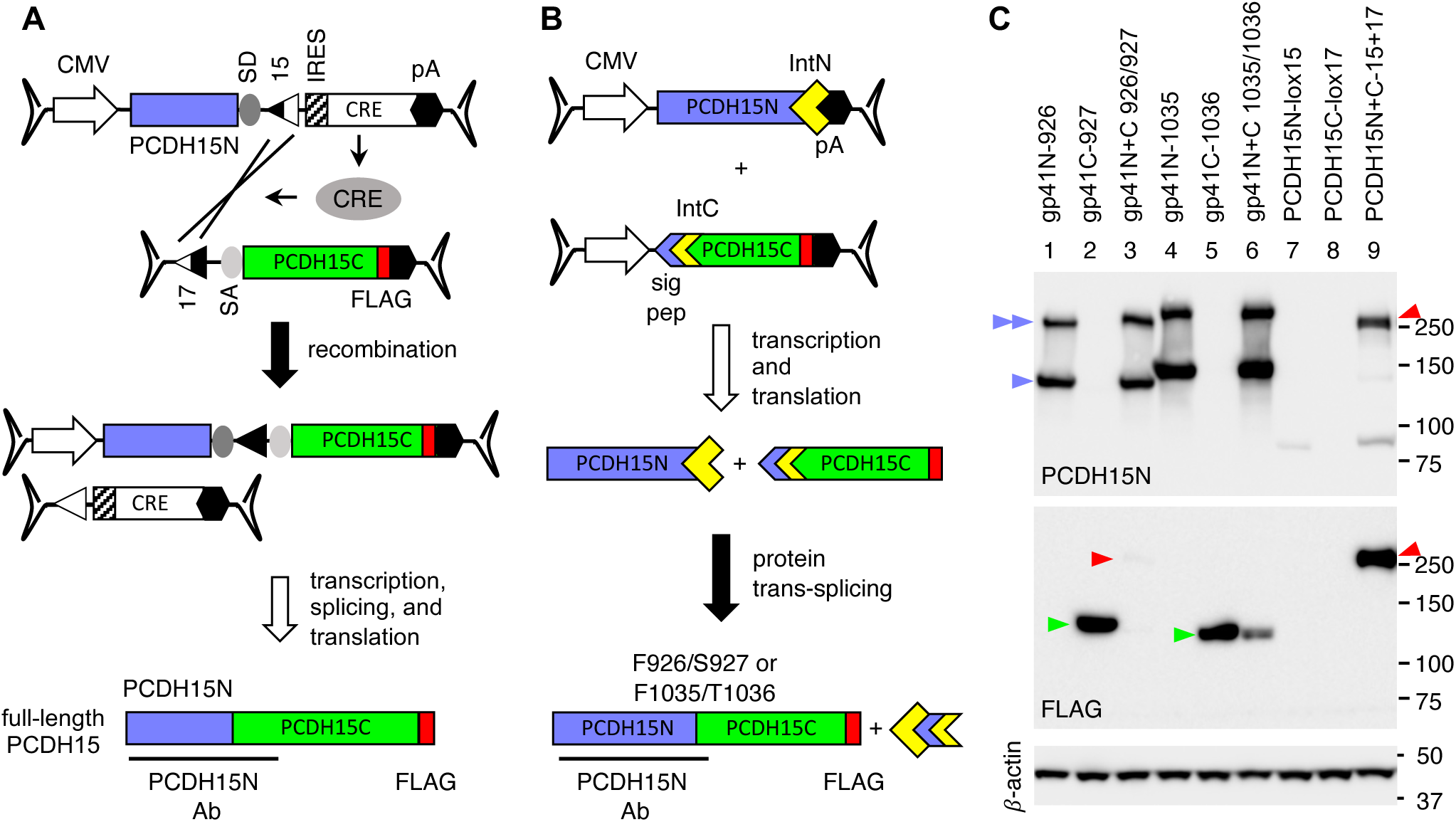
Reconstitution of PCDH15 using the gp41 split intein and the CRE-lox recombination approaches. (A and B) Strategies for the PCDH15 reconstitution using dual AAV-*PCDH15* vectors. *PCDH15* CDS was split at E644/G645 for the CRE-lox set (A) and at F926/S927 or F1035/T1036 for the gp41 sets (B). A signal peptide derived from PCDH15 (N-terminal 26 residues) was added to the N-terminus of IntC to facilitate the extracellular translocation of IntC. PCDH15N antibody was raised against the N-terminal half of human PCDH15 (aaQ27-A1376). A FLAG tag (red) was added to the C-terminus of the protein. IRES: internal ribosome entry site, sig pep: signal peptide for extracellular translocation. 15: loxJT15, 17: loxJTZ17, SD: splice donor site, SA: splice acceptor site, pA: polyA signal. (C) Reconstitution of PCDH15 by the CRE-lox approach. HEK293T cells were transduced with AAV-*PCDH15* vectors as indicated (lanes 1-3: gp41 926/927 set; lanes 4-6: gp41 1035/1036 set; lanes 7-9: CRE-lox set), and cell lysates were subjected to SDS-PAGE and immunoblotting with PCDH15N and FLAG antibodies. The single blue arrowhead marks monomeric forms of PCDH15 N-terminal truncated protein products derived from the gp41N vectors, and the double blue arrowhead indicates presumptive homodimers of PCDH15 N-terminal truncated proteins. The green arrowheads indicate C-terminal truncated proteins from the gp41C vectors. Red arrowheads mark the reconstituted full-length PCDH15 proteins.

For the CRE-lox set, the full-length human *PCDH15* CDS (5,865 bp) was divided into two segments (1,932 bp and 3,933 bp) and inserted into the 5’ and 3’ vectors, respectively (**Figure 6A**). Similar to the *IFT140* gene, the *CRE* CDS was included within the 5’ vector, but in this case, an internal ribosome entry site (IRES) was used instead of the T2A self-cleaving peptide. For the split-intein set, we designed two variations, in which PCDH15 was split at F926/S927 and F1035/T1036 (**Figure 6B**). PCDH15 is a single-pass transmembrane protein, with its N-terminal two-thirds located extracellularly and the remaining portion on the cytoplasmic side. While a signal peptide is present at the N-terminus of PCDH15 (and PCDH15_N) for extracellular translocation, it is absent in the C-terminal half. To facilitate the extracellular translocation of IntC and ensure the presence of IntN and IntC in the same cellular compartments, we introduced the signal peptide of PCDH15 (N-terminal 26 residues) to IntC. Additionally, we introduced a FLAG tag to the C-terminus of PCDH15 in both CRE-lox and split-intein sets to facilitate protein detection. For the detection of the N-terminal portion of PCDH15, a polyclonal antibody (PCDH15N Ab) raised against recombinant human PCDH15 protein (aa Q27-A1376) was used.

When transduced to 293T cells (at an MOI of 3×10^4^ per vector), the CRE-lox-based dual AAV-*PCDH15* vectors demonstrated robust production of full-length PCDH15 (**Figure 6C**, lane 9; red arrowheads). In contrast, although the split-intein-based AAV-*PCDH15* vectors efficiently produced their respective truncated proteins when individually transduced (lanes 1,2,4, and 5; blue and green arrowheads), the reconstituted full-length proteins were either barely detectable or undetectable (lanes 3 and 6). Interestingly, a notable amount of PCDH15 N-terminal truncated proteins was observed in the 250-350 kDa range without the C-terminal half of PCDH15 (lanes 1 and 4; double blue arrowheads). PCDH15 is known to form homodimers through its extracellular domain (50), and we speculate that the high molecular weight products represent homodimers of the PCDH15 N-terminal truncated proteins. Another intriguing observation was the reduction of unconjugated PCDH15 C-terminal truncated protein products (green arrowheads) when co-expressed with PCDH15_N. Although the reason for this phenomenon remains unknown, we speculate that it might be linked to the degradation of reconstituted proteins, possibly attributable to issues related to protein structure or folding. Overall, these results indicate that the CRE-lox method is a more suitable approach for *PCDH15*.

### Reconstitution of CEP290 by CRE-lox-mediated recombination

*CEP290* mutations are associated with various ciliopathies, ranging from isolated retinal dystrophy to syndromic conditions such as Bardet-Biedl syndrome, Joubert syndrome, and Meckel-Gruber syndrome (7–9, 51–53). Although the full-length human *CEP290* CDS (7,440 bp) can be accommodated in two AAV vectors, a better outcome was obtained when *CEP290* was split into three AAV vectors in a prior study to develop *CEP290* gene therapy vectors using the split intein strategy (29). However, even with the optimized set, the yield of full-length CEP290 was very low, and a large amount of truncated protein products remained unspliced.

We applied the CRE-lox approach to *CEP290* and developed bipartite and tripartite AAV-*CEP290* vectors. For the bipartite set (**Figure 7A**), the *CEP290* CDS was divided into two fragments (3,527 bp and 3,913 bp) and inserted into the 5’ and 3’ vectors, respectively. CRE was delivered through a separate AAV vector (AAV-EF1α-*DD*-*CRE*). To prevent constitutive overexpression of *CRE*, we adopted dihydrofolate reductase destabilizing domain (DD)-fused CRE (54, 55), which undergoes rapid degradation through the proteasomal pathway but can be temporarily stabilized by the addition of trimethoprim (TMP) (**Figure S4**). For the tripartite set (**Figure 7C**), *CEP290* was split into 3 segments (5’ (E1): 1,065 bp, middle (E2): 2,913 bp, and 3’ (E3): 3,462 bp), and the *CRE* gene was included within the 5’ vector. The loxJT15 and loxJTZ17 pair was used to join the 5’ and middle vectors and the lox15:2272 and lox17:2272 pair was used for the middle and 3’ vectors. A FLAG tag sequence was introduced at the 5’ end of *CEP290* to facilitate the detection of the N-terminal portion of CEP290 in both sets, and the CEP290-C antibody was used to detect the C-terminal portion of CEP290.

**Figure 7.**
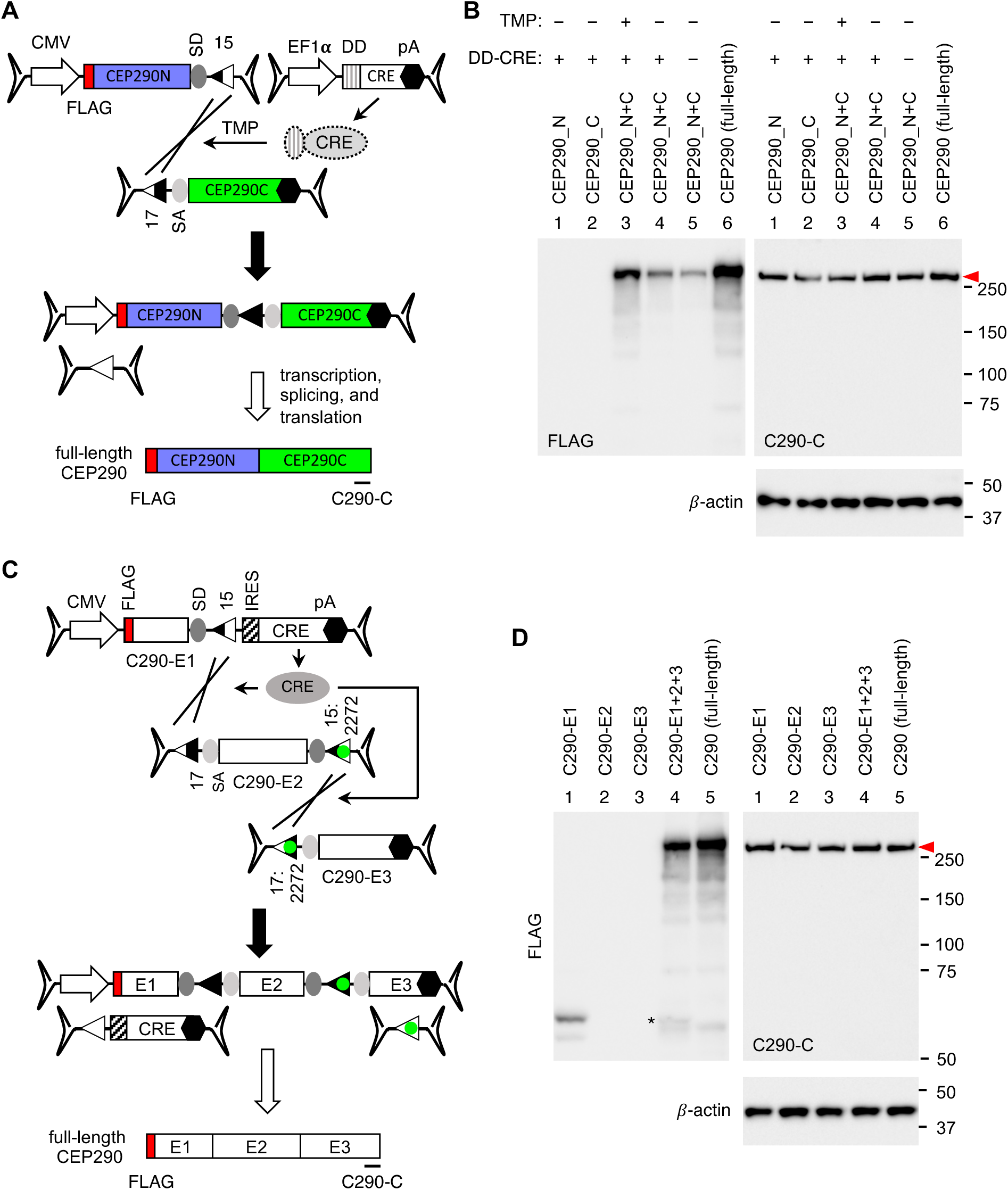
CRE-lox mediated reconstitution of *CEP290*. (A) Schematic representation of the bipartite AAV-*CEP290* vectors. The 5’ vector is composed of a CMV promoter, 5’ 3,527 bp of human *CEP290*, an SD site, and a loxJT15 site. The 3’ vector consists of a loxJTZ17 site, an SA site, 3’ 3,913 bp of *CEP290*, and a BGH polyA signal. A FLAG tag was added to the 5’ end of *CEP290*. Destabilizing domain (DD)-fused CRE was delivered via a separate AAV vector (AAV-EF1α-*DD-CRE*). The location of the C290-C antibody epitope was marked by a black line at the bottom of the schematic. (B) Production of full-length CEP290 proteins (red arrowhead) by CRE-lox-mediated recombination in 293T cells. HEK293T cells were transduced with dual AAV2/2-*CEP290* (MOI: 3×10^4^ per vector) and AAV2/2-EF1α-*DD-CRE* (MOI: 1×10^4^) vectors. After transduction, cells were treated with 10 μM trimethoprim (TMP) for 48 hours to stabilize DD-CRE, and cell lysates were subjected to SDS-PAGE followed by immunoblotting with FLAG and C290-C antibodies. Lane 1: AAV-CEP290_N only, lane 2: AAV-CEP290_C only, lanes 3-4: dual AAV-CEP290_N+C, and lane 5: pSS-FS-hCEP290 plasmid transfected (full-length; positive control). Cells in lanes 1-3 were co-transduced with AAV2/2-EF1α-*DD-CRE*. Numbers on the right denote the location of protein standards. β-actin was used as a loading control. (C) Schematic representation of the tripartite AAV-*CEP290* vectors. The 5’ vector (E1) is composed of a CMV promoter, 5’ 1,065 bp of human *CEP290*, an SD site, a loxJT15 site, IRES, *CRE* CDS, and a BGH polyA signal. The middle vector (E2) is composed of a loxJTZ17 site, an SA site, 2,913 bp of *CEP290* CDS, an SD site, and a lox15:2272 site. The 3’ vector (E3) consists of a lox17:2272 site, an SA site, 3’ 3,462 bp of *CEP290*, and a BGH polyA signal. Others are the same as in panel A. (D) Production of full-length CEP290 proteins (red arrowhead) by CRE-lox-mediated recombination in 293T cells. Others are the same as in panel B.

To evaluate the reconstitution and expression of *CEP290* from the bipartite AAV-*CEP290* vectors, HEK293T cells were transduced with the dual AAV-*CEP290* vectors at an MOI of 3×10^4^ (per vector) and AAV-EF1α-*DD-CRE* at an MOI of 1×10^4^. After transduction, cells were treated with 10 μM TMP for 48 hours to stabilize DD-CRE, and *CEP290* expression was assessed by immunoblotting (**Figure 7B**). FLAG-tagged, full-length CEP290 expression was detectable in cells transduced with all three AAV vectors and treated with TMP (lane 3). FLAG-CEP290 expression was also detectable without TMP treatment (lane 4), but at a significantly lower level (average 24% of TMP-treated cells; n=3). A small amount of FLAG-CEP290 was observed in the absence of CRE (lane 5; average 9% of TMP-treated cells; n=3), representing *CEP290* gene reconstitution via spontaneous recombination between the dual AAV-*CEP290* vectors. Plasmid DNAs encoding full-length *CEP290* were transfected and used as a positive control (lane 6). For some unknown reason, over-expressed CEP290 appears to be unstable and degraded, and consequently, we were not able to observe increased CEP290 protein levels beyond endogenous expression using the CEP290-C antibody. β-actin was used as a loading control.

The tripartite AAV-*CEP290* vectors were transduced into 293T cells at an MOI of 3×10^4^ (per vector), and FLAG-CEP290 expression was examined as above. As shown in **Figure 7D**, full-length *CEP290* expression was detectable only when all three AAV vectors were transduced. These data demonstrate that both bipartite (with a separate AAV-*CRE* vector) and tripartite AAV-*CEP290* vectors efficiently deliver the full-length *CEP290* gene to target cells.

### Reconstitution of CDH23 by CRE-lox mediated recombination

We then applied the CRE-lox-mediated DNA recombination approach to *CDH23* gene therapy vectors. Inactivating mutations of *CDH23* cause Usher syndrome type 1D (USH1D) (12, 13). The full-length human *CDH23* CDS spans 10,065 bp, requiring three AAV vectors for delivery. Since the total payload capacity of three AAV vectors is ∼14 kb (excluding ITRs), we split the *CDH23* CDS into 3 fragments (5’ vector (E1): 2,176 bp, middle vector (E2): 4,077 bp, and 3’ vector (E3): 3,812 bp) and included the *CRE* gene (with a T2A peptide) in the 5’ vector (**Figure 8A**). The loxJT15 and loxJTZ17 pair was used to join the 5’ and the middle vectors and the lox15:2272 and lox17:2272 pair was used for the middle and the 3’ vectors. CDH23 is a type-I single transmembrane protein with an N-terminal signal peptide, and an HA tag was inserted after the signal peptide for protein detection. A CBh promoter and a BGH polyA signal were used as a promoter and a transcription termination signal, respectively.

**Figure 8.**
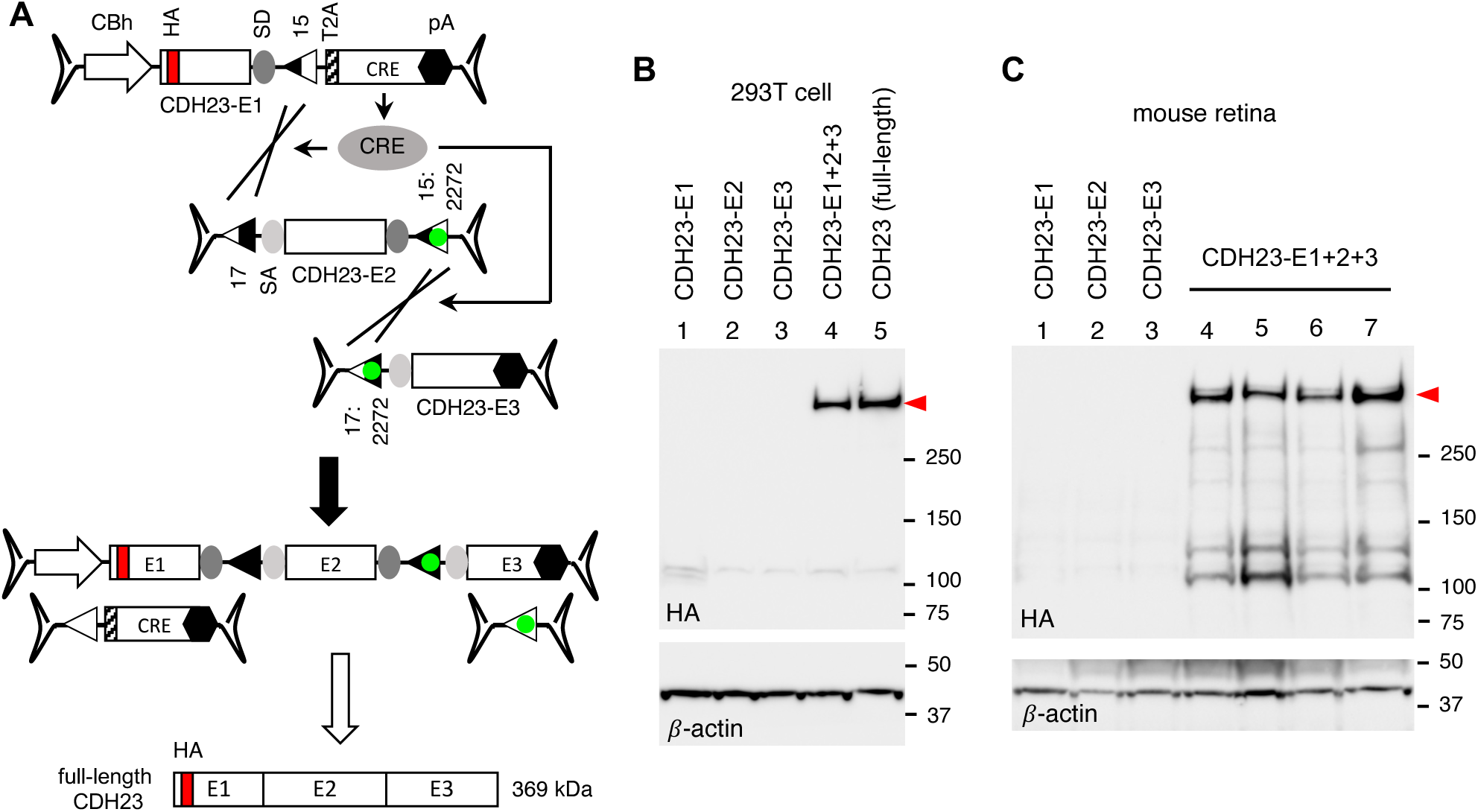
CRE-lox-mediated reconstitution of *CDH23* delivered by tripartite AAV vectors. (A) Schematic of the *CDH23* reconstitution using tripartite AAV-*CDH23* vectors. The *CDH23* CDS (10,065 bp) was split into three pieces (E1: 2,176 bp, E2: 4,077 bp, and E3: 3,812 bp), and the *CRE* gene was included in the 5’ (E1) vector for self-inactivation after recombination. A T2A “self-cleaving” peptide was used for *CRE* expression. An HA tag (red) was added to the N-terminus of CDH23 for detection (after the signal peptide). (B) Production of full-length CDH23 proteins (red arrowhead) by CRE-lox-mediated recombination in 293T cells. HEK293T cells were transduced with tripartite AAV2/2-*CDH23* vectors (MOI: 3×10^4^ per vector), and cell lysates were subjected to SDS-PAGE followed by immunoblotting with HA antibodies. Lane 1: AAV-CDH23-E1 only, lane 2: AAV-CDH23-E2 only, lane 3: AAV-CDH23-E3 only, lane 4: AAV-CDH23-E1, E2, and E3 co-transduced, and lane 5: pSS-HA-CDH23-SF plasmid transfected (full-length; positive control). β-actin was used as a loading control. (C) Expression of *CDH23* from the tripartite AAV-*CDH23* vectors in mouse retinas. Tripartite AAV-*CDH23* vectors were subretinally administered to wild-type mice as indicated (lane 1: E1 vector alone, lane 2: E2 vector alone, lane 3: E3 vector alone, lanes 4-7: E1, E2, and E3 co-injected) at the dose of 3×10^9^ GC per vector. Treated eyes were collected 3 weeks post-injection and retinal protein extracts were subjected to SDS-PAGE and immunoblotting. Each lane represents individual eyes.

To test whether the tripartite AAV-*CDH23* vectors could deliver full-length *CDH23*, we transduced HEK293T cells with these vectors at an MOI of 3×10^4^ (per vector; serotype AAV2) and examined the production of full-length CDH23 proteins by immunoblotting. When all three vectors were co-transduced, robust expression of *CDH23* was observed (**Figure 8B**; red arrowhead). A plasmid containing a full-length *CDH23* expression cassette was used as a positive control (lane 5), and β-actin was used as a loading control. To test whether these vectors could be used to deliver *CDH23 in vivo*, we administered the tripartite AAV-*CDH23* vectors to wild-type mouse eyes by subretinal injections (serotype: AAV5, dose: 3×10^9^ GC per vector). Consistent with the results in 293T cells, full-length CDH23 proteins were readily detected in all eyes injected with the tripartite AAV vectors (**Figure 8C**, lanes 4-7).

## Discussion

Reconstitution of therapeutic genes via the CRE-lox-mediated DNA recombination offers several advantages compared to other approaches that have been reported thus far.

First, compared to natural recombination-dependent approaches such as trans-splicing, overlapping, and hybrid dual or triple AAV approaches (16–19), the CRE-lox-mediated recombination drastically improves the recombination efficiency and the yield of correctly reconstituted genes. This is especially true when tripartite or quadripartite AAV vectors are used. The use of non-compatible, hybrid lox sites prevents the excision of floxed sequences, ensures recombination in a pre-determined configuration, and inhibits the disassembly of reconstituted genes (**Figure S1D-F**). The improved efficiency and yield reduce the number of AAV particles needed to transduce target cells, and the use of fewer AAV vectors reduces the potential risks of viral vector-derived toxicity and inflammation.

Second, the CRE-lox-mediated DNA recombination approach provides more flexibility regarding splitting positions compared to the protein trans-splicing approach. The efficiency of protein trans-splicing is influenced by the amino acid residues adjacent to split inteins (24, 30, 31). The first residue within the C-extein is particularly important, and Cys, Ser, and Thr residues are strongly preferred. This constraint greatly limits the number of possible locations where a protein may be split. Moreover, protein truncations can affect a protein’s structure, stability, and localization, and these factors also influence the overall efficiency and yield of the protein reconstitution. If the target gene encodes a transmembrane or secreted protein (e.g., PCDH15 and CDH23), the topology and secretion of each protein fragment should be considered when determining splitting positions. Ideal splitting positions for the split intein approach should support the protein trans-splicing process and have minimal or no effect on the folding and stability of each truncated protein fragment and reconstituted proteins. Additionally, the truncated protein products should be localized to the same compartment or close locations to increase the likelihood of engagement, while the size of each fragment should be small enough to fit into single AAV vectors. Therefore, identifying optimal splitting positions could be challenging and usually involves comparing multiple candidate sites empirically. And the complexity increases if the target gene requires 3 or 4 AAV vectors. In contrast, when using the CRE-lox approach, protein structure, stability, localization, and topology are not factors to consider since the reconstitution occurs at the DNA level. Cargo capacity expansion can be achieved by simply adding additional sets of non-compatible lox sites to AAV vectors.

Another significant advantage of the CRE-lox approach over protein trans-splicing is the lack or minimal production of truncated proteins. Protein trans-splicing requires the production of “half” proteins before reconstitution, which can have dominant negative or harmful effects if continuously expressed. In contrast, truncated protein production is either absent or low when the CRE-lox strategy is used because the AAV vectors lack a promoter or a polyA signal, which stabilizes mRNA. Although ITRs have some intrinsic promoter activities (56), they are weak in most cells. When the *CRE* gene is included in the 5’ vector, a truncated protein and CRE are initially produced. However, as recombination progresses and the 5’ vector is converted to a full-length therapeutic cassette, the production of truncated proteins and CRE diminishes. The lack or minimal production of truncated proteins may be crucial for certain genes if such protein products are toxic to cells, and the “self-inactivation” feature of the CRE-containing 5’ vector provides an additional layer of safety to the CRE-lox approach.

Lastly, although the CRE-lox approach requires the delivery of CRE in addition to therapeutic genes, the practical payload capacity of AAV vectors with the CRE-lox approach is either comparable with or larger than that of the split intein-based approach. The protein trans-splicing approach requires each AAV vector to have its own promoter and transcription termination signal to produce therapeutic gene products and split inteins. The repeated inclusion of transcriptional regulatory elements erodes the AAV vector’s combined payload capacity. In contrast, the CRE-lox approach only requires one promoter and one transcriptional termination signal for the entire vector set, which becomes more beneficial as more AAV vectors are needed.

One notable concern of the CRE-lox approach is the prolonged expression of CRE in transduced cells. Prolonged expression of *CRE* could lead to unintended recombination in the human genome, potentially resulting in unwanted mutations or genomic instability. Moreover, prolonged *CRE* expression could also lead to immune responses, which may limit the effectiveness of AAV gene therapies. This could potentially result in the destruction of cells expressing the therapeutic gene or a reduction in the efficacy of the AAV gene therapy over time. In this regard, the inclusion of *CRE* in 5’ vectors and being “self-inactivated” by recombination significantly reduces these risks. Alternatively, an inducible promoter or destabilizing domain-fused CRE may be used to control *CRE* expression. Other site-specific DNA recombinases that display higher specificity than CRE may be used as well. It is noteworthy that while numerous transgenic rodent lines expressing *CRE* have been created and some phenotypes have been reported (57), constitutive expression of *CRE* does not appear to cause serious health concerns in rodents. However, a thorough investigation of genes or sites in the human genome that could be modified by CRE will be necessary. As with any medical treatment, it is crucial to carefully consider the potential risks and benefits associated with CRE-lox-dependent AAV gene therapies.

In summary, the CRE-lox approach described in this study offers a simple, versatile, and efficient platform for producing AAV-based, generic gene replacement therapy vectors capable of delivering large genes. As this approach delivers full-length genes, the gene therapy vectors developed using this method have the potential to be generally applicable to all patients with loss-of-function mutations.

## Materials and Methods

### Plasmid DNAs

To generate the GFP expression reporter loxP-N (**Figure 2**), a double-stranded DNA fragment corresponding to the loxJT15+C290C+17:m7+FLAG+17:HT1+HA+17:HT2+MYC+17:N+V5 portion was synthesized (Integrated DNA Technologies) and inserted at the C-terminus of GFP within the pEGFP-C3 plasmid (Clontech) using a GenBuilder Cloning kit (GenScript). Reporter loxP-2272 was generated by substituting the lox17:N site (ATAACTTCGTATA<u>GTATACCT</u>TATAGCAATTTAT) within loxP-N with the lox17:2272 site (ATAACTTCGTATA<u>GGATACTT</u>TATAGCAATTTAT) using a Q5 Site-Directed Mutagenesis kit (New England Biolabs) and PCR. Reporter lox2272-N was produced by replacing the loxJT15 site (AATTATTCGTATA<u>GCATACAT</u>TATACGAAGTTAT) in the loxP-N reporter with the lox15:2272 site (AATTATTCGTATA<u>GGATACTT</u>TATACGAAGTTAT). Reporters loxP-HT2 and lox2272-HT2 were derived by eliminating the lox17:N sites in loxP-N and lox2272-N using a Q5 Site-Directed Mutagenesis kit and PCR. The DNA sequences downstream of GFP are shown in the **Supplementary Material**.

The AAV shuttle plasmids used in this study are listed in the **Supplementary Material**, and their sequences are available upon request. Briefly, the pFBAAV-related plasmids were generated by modifying the pFBAAVCMVmcsBGHpA plasmid (from the Viral Vector Core Facility, University of Iowa). Plasmids pAAV-CMV-gp41-IntC and pAAV-CBh-JT1522 were custom-synthesized and procured from VectorBuilder. Other pAAV-related plasmids were developed by modifying these plasmids using standard molecular biology techniques and the GenBuilder Cloning kit (GenScript). PCR primers and oligonucleotides used were obtained from Integrated DNA Technologies (IDT). Plasmids pcDNA3-MYC-hBBS1, pCS2FLAG-hLZTFL1, and pCS2HA-hLZTFL1 were previously described (58, 59). Plasmid DNAs containing human *IFT140* (BC035577) and *PCDH15* (NM_033056.4) coding sequences (CDS) were acquired from transOMIC and GenScript, respectively. These plasmids were used as PCR templates to produce pFBAAV-CMV-IFT140N-15, pAAV-CMV-IFT140N-1522-CRE, pAAV-CBh-IFT140N-1522-CRE, pFBAAV-1722-IFT140C-pA, pAAV-CMV-IFT140N-gp41N, pAAV-CMV-gp41C-IFT140C, pAAV-CMV-PCDH15N-926, pAAV-CMV-gp41C-PCDH15C-927, pAAV-CMV-PCDH15N-1035, pAAV-CMV-gp41C-PCDH15C-1036, pAAV-CMV-PCDH15N-15-IRES-CRE, and pFBAAV-17-PCDH15C-pA. A plasmid containing the full-length human *CEP290* CDS (NM_025114.4) was reported earlier (60) and used as a PCR template for bipartite (pFBAAV-CMV-CEP290N-JT15 and pFBAAV-17-CEP290C-pA) and tripartite AAV-*CEP290* vectors (pAAV-CBh-CEP290-E1-15-CRE, pAAV-17-CEP290-E2-1522, and pFBAAV-1722-CEP290-E3-pA). The human *CDH23* CDS was divided into 5 segments and synthesized by IDT according to the NM_022124.6 sequence. An HA tag was introduced after CDH23’s signal peptide (N-terminal 28 residues). These 5 segments were PCR-amplified using Q5 High-Fidelity DNA polymerase and inserted into the multiple cloning site (MCS) of pAAV-CBh-JT15-CREv2 (first segment), pAAV-JTZ17-mid-1522 (second and third segments), and pFBAAV-1722-pA (fourth and fifth segments) using the GenBuilder Cloning kit to produce pAAV-CBh-CDH23-E1-15-CRE, pAAV-17-CDH23-E2-1522, and pFBAAV-1722-CDH23-E3-pA, respectively. A full-length *CDH23* expression vector, pSS-HA-hCDH23-SF, was generated by PCR amplification of the CDH23-E1, E2, and E3 fragments and insertion of these 3 fragments into the MCS of the pSS-SF plasmid (61) using the GenBuilder Cloning kit (sequence available upon request).

### Transfection and AAV transduction in HEK293T/17 cells

HEK293T/17 cells were acquired from American Type Culture Collection (ATCC; #CRL-11268) and cultured in Dulbecco’s Modified Eagle’s Medium (DMEM; Gibco) supplemented with 10% (v/v) fetal bovine serum (FBS; Gibco) and 100 units/ml penicillin/streptomycin (Gibco) at 37 °C in a humidified 5% CO2 incubator. Plasmid DNAs were transfected in 12-well plates (Sarstedt) using FuGENE HD Transfection Reagent (Promega) following the manufacturer’s instructions. Twenty-four hours post-transfection, cells were transferred to 6-well plates and cultured for 48 hours before harvesting.

AAV vectors with serotype 2 were produced by VectorBuilder and used for transducing HEK293T/17 cells. Approximately 5×10^4^ cells/well were seeded in a 24-well plate (Sarstedt) 24 hours prior to transduction. On the day of transduction, AAV vectors were thawed on ice, and appropriate volumes were taken to achieve the desired MOI and added to 250 μl of DMEM with 2% FBS. After removing the existing culture medium, the AAV-containing medium was added to the cells. The cells were then incubated for 16-17 hours in the cell culture incubator. Transduced cells were expanded in 6-well plates with complete culture medium and further incubated for 72 hours before harvesting. For trimethoprim (TMP) treatment, transduced cells were expanded in 6-well plates with complete medium containing 10 μM TMP (Sigma), cultured for 48 hours, and then switched to a normal culture medium for an additional 24 hours before harvesting.

### Mouse, animal study approval, and AAV subretinal injection

All animal procedures were approved by the Institutional Animal Care and Use Committee (IACUC) of the University of Iowa and conducted following the recommendations in the Guide for the Care and Use of Laboratory Animals of the National Institutes of Health. All animals were maintained in 12-hour light-dark cycles and fed standard mouse chow *ad libitum*.

For subretinal injections, AAV vectors with serotype 5 (VectorBuilder) were used. On the day of injection, AAV2/5 vectors were thawed on ice and diluted to desired titers (2.5×10^9^ GC/μl per vector for AAV-IFT140 and 1.5×10^9^ GC/μl per vector for AAV-CDH23) in PBS supplemented with 0.001% Poloxamer 188 (Sigma Aldrich).

One-to-two-month-old wild-type C57BL/6J mice (both males and females) were used for AAV subretinal injections. Mice were anesthetized with a ketamine/xylazine mixture (87.5 mg/kg ketamine, 12.5 mg/kg xylazine), and 10% povidone-iodine and 1% tropicamide solutions were applied. Under a Zeiss OPMI VISU 150 surgical microscope, eyes were gently rotated using forceps, and a conjunctival peritomy was made with Vannas scissors, followed by a sclerotomy made posterior to the limbus using a 30-gauge needle. Transscleral subretinal injections were performed with a Hamilton syringe attached to a blunt-end 32-gauge needle inserted subretinally at an oblique angle, and 2 μl of vector solutions were delivered. Antibiotic/steroid ophthalmic ointment (neomycin and dexamethasone) was applied after injections. If significant hemorrhage was observed, the animal was excluded from follow-up analysis.

### Protein extraction, SDS-PAGE, and immunoblotting

To extract proteins from HEK293T/17 cells, cells in 6-well plates were briefly rinsed with ice-cold PBS and lysed with 300 μl of ice-cold lysis buffer (50 mM HEPES pH 7.0, 150 mM NaCl, 2 mM EGTA, 2 mM MgCl2, 1% Triton X-100) supplemented with Protease Inhibitor Cocktail (Bimake). Lysates were centrifuged at 20,000x g for 15 minutes at 4°C to precipitate insoluble materials.

To extract proteins from mouse retinas, mice were euthanized by CO2 asphyxiation followed by cervical dislocation. Eyes were enucleated and anterior segments were removed using micro-dissecting scissors. Collected tissues were homogenized with a pestle in a lysis buffer (50 mM HEPES pH 7.0, 150 mM NaCl, 2 mM EGTA, 2 mM MgCl2, 1% Triton X-100) supplemented with the Protease Inhibitor Cocktail. Homogenates were centrifuged at 20,000x g for 15 minutes at 4°C to precipitate insoluble materials.

Supernatants were mixed with NuPAGE LDS Sample Buffer (Invitrogen) and Reducing Agent (Invitrogen), and 40-50 μg of proteins were loaded per lane on NuPAGE 4-12% (wt/vol) Bis-Tris gels (Invitrogen; for all genes except *CDH23*) or 3-8% Tris-Acetate gels (Invitrogen; for *CDH23*). Proteins were transferred onto nitrocellulose membranes (BioRad) and immunoblotting was performed following standard protocols. Antibodies used are listed in **Table S1**. Proteins were detected by using horse radish peroxidase (HRP)-conjugated secondary antibodies and SuperSignal West Dura Extended Duration Substrate (Thermo Scientific). Images were taken with a ChemiDoc Imaging system (Bio-Rad).

## Acknowledgments

This work was supported by National Institutes of Health grant R01-EY034176 (to S.S.), Retina Research Foundation Pilot grant (to S.S.), Flagella Vision Foundation (to S.S. and A.V.D.), and the Ronald Keech professorship (to A.V.D.). This work was also supported in part by National Institutes of Health grant R30-EY025580 to the University of Iowa.

## Author contributions

Conception and study design: S.S., vector design: S.S., materials: P.D., K.D.R., R.J.S, subretinal injection: P.D., K.D.R., J.M.T., Y.H., S.H., A.V.D., data collection: P.D., K.D.R., S.S., data analysis: S.S., funding acquisition: S.S., A.V.D., manuscript drafting: S.S., manuscript revision: K.D.R., Y.H., and A.V.D.

## Abbreviations

AAV: adeno-associated virus
BGH: bovine growth hormone
CDS: coding sequence
DD: destabilizing domain
GC: genome copies
HRP: horse radish peroxidase
IntC: C-terminal split intein
IntN: N-terminal split intein
IRES: internal ribosome entry site
ITR: inverted terminal repeat
LCA: Leber congenital amaurosis
LE: left element
MOI: multiplicity of infection
RE: right element
RP: retinitis pigmentosa
SA: splice acceptor
SD: splice donor
TMP: trimethoprim

