## Supplementary Material for "Delivering large genes using adeno-associated virus and the CRE-lox DNA recombination system"

**The PDF file includes:**

Figure S1. Advantages of the non-compatible, reaction-equilibrium modifying lox sites.

Figure S2. Design of the lox site compatibility reporters.

Figure S3. Schematic of the IFT140 domain organization and the selected splitting position for the split intein approach.

Figure S4. Stabilization of DD-CRE by trimethoprim (TMP).

Plasmids used in this study.

1) DNA sequences of the lox site incompatibility reporters (downstream of GFP)

2) AAV shuttle vectors

3) Other expression plasmids used

Table S1. Antibodies used in this study.

Additional References

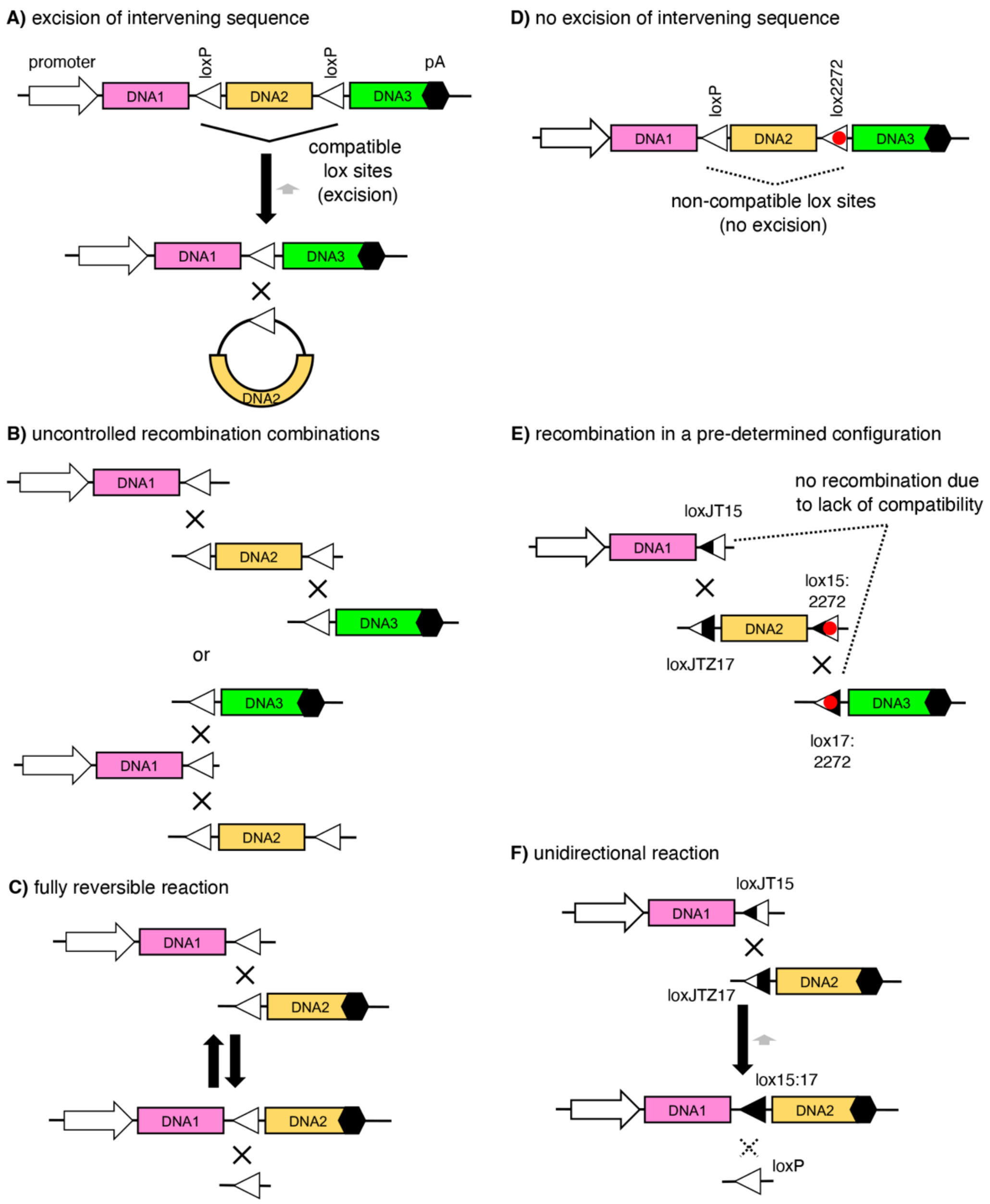

**Figure S1**. Advantages of the non-compatible, reaction-equilibrium modifying lox sites.
(A-C) Problems of using only one species of or compatible lox sites to splice more than two DNA molecules. (A) The presence of multiple compatible lox sites within a single DNA molecule leads to a rapid excision of intervening sequences. (B) Since all DNA fragments possess compatible lox sites, recombination reactions can occur in various combinations, including unintended ones. (C) Because both the substrates and the products have compatible lox sites, the recombination reaction becomes fully reversible, causing the reconstituted DNAs to continuously cycle through assembly-disassembly with a 50:50 equilibrium ratio.
(D-F) We have devised 5 pairs of novel lox sites by combining non-compatible (loxm7, loxN, lox2272, loxHT1, and loxHT2) and reaction-equilibrium modifying lox sites (loxJT15 and loxJTZ17). (D) Non-compatible lox sites (e.g., loxP and lox2272) prevent the excision of intervening sequences. (E) By employing two or more non-compatible lox sites, one can specify the DNA molecules to recombine. (F) Incorporating reaction-equilibrium modifying lox sites (e.g., loxJT15 and loxJTZ17) enhances the yield of reconstituted DNAs by preventing the disassembly of the reconstituted expression cassettes.

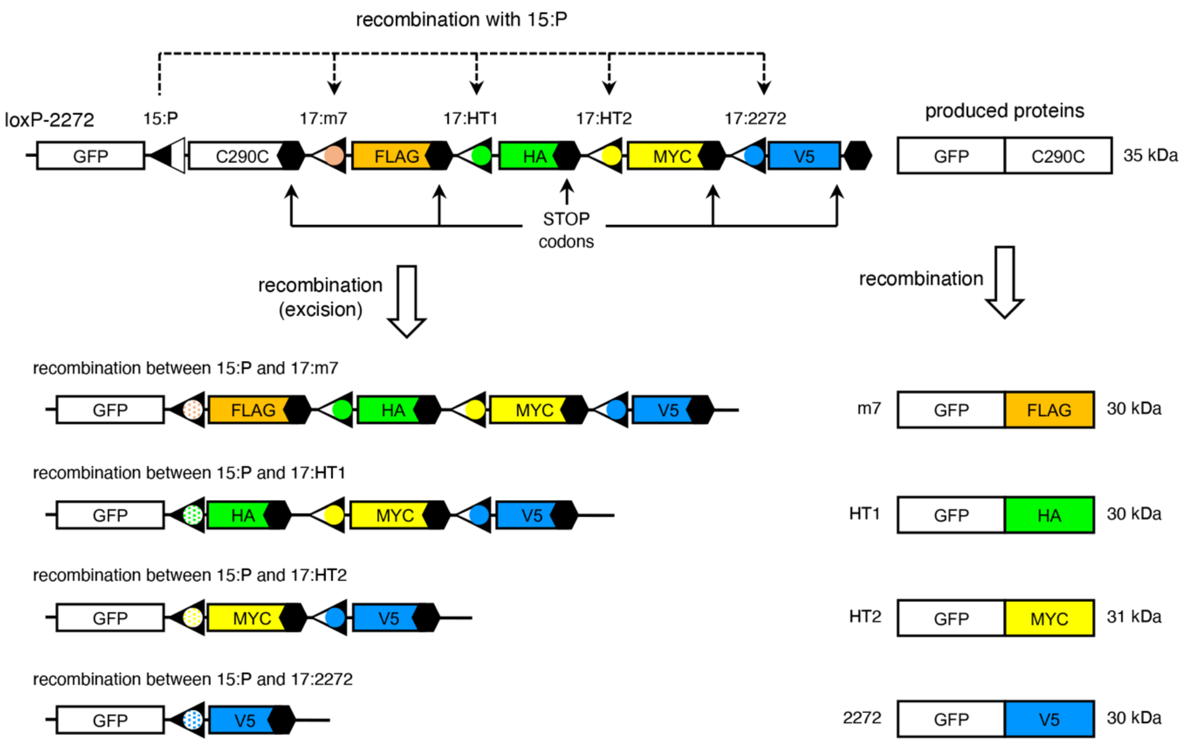

**Figure S2**. Design of the lox site compatibility reporters.
Schematics of loxP-2272 are shown as a representative. In the absence of recombination, GFP+C290C fusion proteins (~35 kDa) are produced, and translation stops at the end of C290C due to the presence of a STOP codon. If recombination takes place between 15:P and 17:m7 sites, the C290C portion is excised and the FLAG tag is linked to GFP in-frame, resulting in the production of GFP+FLAG fusion proteins (30 kDa). If recombination takes place between 15:P and 17:HT1, GFP+HA fusion proteins (30 kDa) are produced. Likewise, recombination of 15:P with 17:HT2 and 17:2272 leads to the production of GFP+MYC and GFP+V5 fusion proteins, respectively.

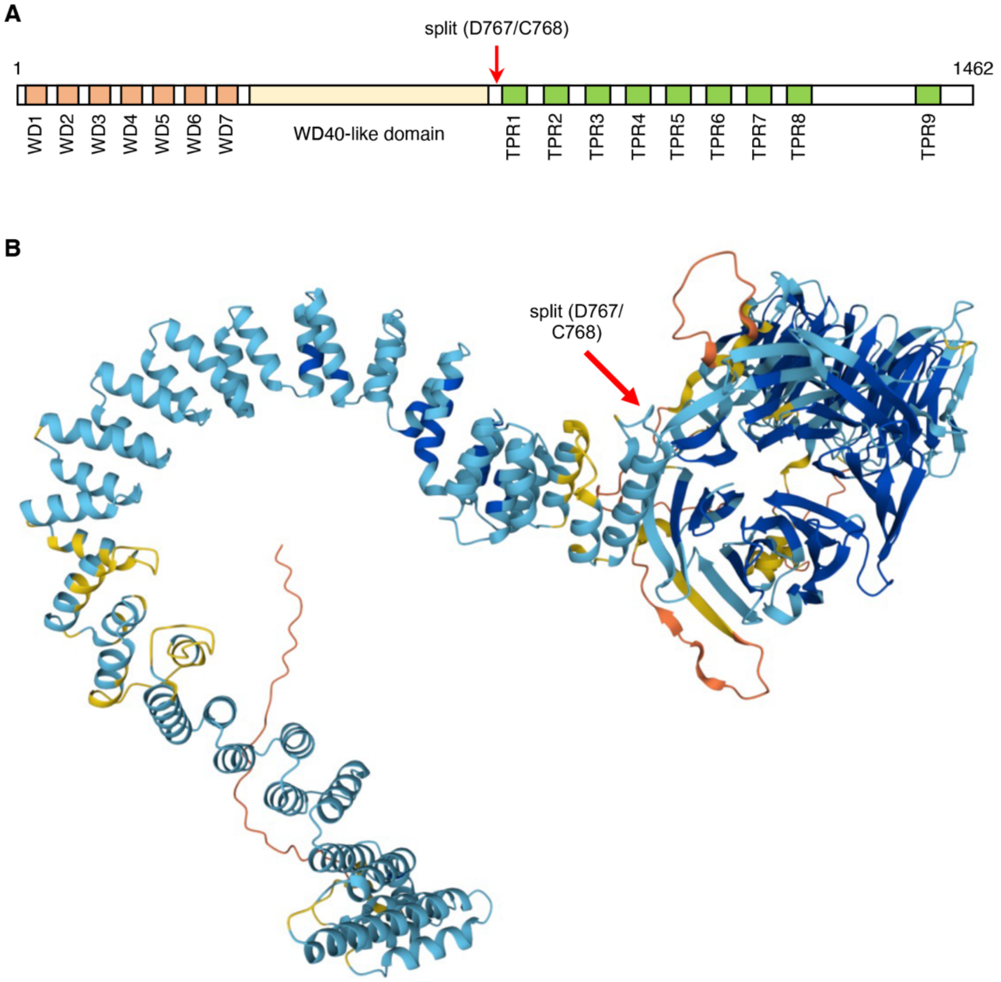

**Figure S3**. Schematic of the IFT140 domain organization and the selected splitting position for the split intein approach.
(A) IFT140 domain organization. The N-terminal portion of IFT140 features 7-blade WD40 repeats, while its C-terminal half encompasses nine tetratricopeptide repeats (TPR).
(B) IFT140 structural model predicted by AlphaFold (identifier: AF-Q96RY7-F1). The model proposes the presence of a WD40-like domain situated in the latter part of the N-terminal half. The splitting position (D767/C768) for the gp41 split intein approach is indicated by the red arrow.

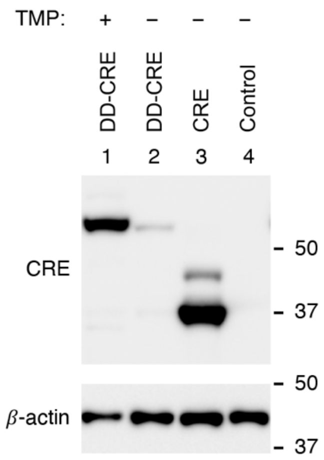

**Figure S4**. Stabilization of DD-CRE by trimethoprim (TMP).
HEK293T cells were transfected with pAAV-EF1α-DD-CRE or pAAV-EF1α-CRE and cell lysates were subjected to SDS-PAGE and immunoblotting. Cells in lane 1 were treated with 10 μM TMP for 48 hours before harvesting. Untransfected cells were used as a negative control and β-actin was used as a loading control.

**Plasmids used in this study**

**1) DNA sequences of the lox site incompatibility reporters (downstream of GFP)**

hybrid lox sites (underlined), C290C (yellow), FLAG (green), HA (cyan), MYC (magenta), and V5 (grey)

loxP-N

tactcaaagcttAATTATTCGTATAGCATACATTATACGAAGTTATgcggccgcAAGTATAATTACAAG
GAAGAAGTGAAGAAGAATATTCTCTTAGAAGAGAAGGTAAAAAAACTTTCAGAACAATTGG
GAGTTGAATTAACTAGCCCTGTTGCTGCTTCTGAAGAGTTTGAAGATGAAGAAGAAAGTCC
TGTTAATTTCCCCATTTACtgATAACTTCGTATATTCTATCTTATAGCAATTTATcggacGACTAC
AAAGACGATGACGACAAGtaaATAACTTCGTATACTATAGCCTATAGCAATTTATatggcTACC
CATACGATGTTCCAGATTACGCTtaATAACTTCGTATATACTATACTATAGCAATTTATtaGAA
CAAAAACTCATCTCAGAAGAGGATCTGtaATAACTTCGTATAGTATACCTTATAGCAATTTATt
ctgtaccGGCAAGCCCATCCCCAACCCCCTGCTGGGCCTGGACAGCACCTAA

loxP-2272

tactcaaagcttAATTATTCGTATAGCATACATTATACGAAGTTATgcggccgcAAGTATAATTACAAG
GAAGAAGTGAAGAAGAATATTCTCTTAGAAGAGAAGGTAAAAAAACTTTCAGAACAATTGG
GAGTTGAATTAACTAGCCCTGTTGCTGCTTCTGAAGAGTTTGAAGATGAAGAAGAAAGTCC
TGTTAATTTCCCCATTTACtgATAACTTCGTATATTCTATCTTATAGCAATTTATcggacGACTAC
AAAGACGATGACGACAAGtaaATAACTTCGTATACTATAGCCTATAGCAATTTATatggcTACC
CATACGATGTTCCAGATTACGCTtaATAACTTCGTATATACTATACTATAGCAATTTATtaGAA
CAAAAACTCATCTCAGAAGAGGATCTGtaATAACTTCGTATAGGATACTTTATAGCAATTTATt
ctgtaccGGCAAGCCCATCCCCAACCCCCTGCTGGGCCTGGACAGCACCTAA

lox2272-N

tactcaaagcttAATTATTCGTATAGGATACTTTATACGAAGTTATgcggccgcAAGTATAATTACAAG
GAAGAAGTGAAGAAGAATATTCTCTTAGAAGAGAAGGTAAAAAAACTTTCAGAACAATTGG
GAGTTGAATTAACTAGCCCTGTTGCTGCTTCTGAAGAGTTTGAAGATGAAGAAGAAAGTCC
TGTTAATTTCCCCATTTACtgATAACTTCGTATATTCTATCTTATAGCAATTTATcggacGACTAC
AAAGACGATGACGACAAGtaaATAACTTCGTATACTATAGCCTATAGCAATTTATatggcTACC
CATACGATGTTCCAGATTACGCTtaATAACTTCGTATATACTATACTATAGCAATTTATtaGAA
CAAAAACTCATCTCAGAAGAGGATCTGtaATAACTTCGTATAGTATACCTTATAGCAATTTATt
ctgtaccGGCAAGCCCATCCCCAACCCCCTGCTGGGCCTGGACAGCACCTAA

loxP-HT2

tactcaaagcttAATTATTCGTATAGCATACATTATACGAAGTTATgcggccgcAAGTATAATTACAAG
GAAGAAGTGAAGAAGAATATTCTCTTAGAAGAGAAGGTAAAAAAACTTTCAGAACAATTGG
GAGTTGAATTAACTAGCCCTGTTGCTGCTTCTGAAGAGTTTGAAGATGAAGAAGAAAGTCC
TGTTAATTTCCCCATTTACtgATAACTTCGTATATTCTATCTTATAGCAATTTATcggacGACTAC
AAAGACGATGACGACAAGtaaATAACTTCGTATACTATAGCCTATAGCAATTTATatggcTACC
CATACGATGTTCCAGATTACGCTtaATAACTTCGTATATACTATACTATAGCAATTTATtaGAA
CAAAAACTCATCTCAGAAGAGGATCTGtaa

lox2272-HT2

tactcaaagcttAATTATTCGTATAGGATACTTTATACGAAGTTATgcggccgcAAGTATAATTACAAGGA
AGAAGTGAAGAAGAATATTCTCTTAGAAGAGAAGGTAAAAAAACTTTCAGAACAATTGGGAG
TTGAATTAACTAGCCCTGTTGCTGCTTCTGAAGAGTTTGAAGATGAAGAAGAAAGTCCTGTT
AATTTCCCCATTTACtgATAACTTCGTATATTCTATCTTATAGCAATTTATcggacGACTACAAAGA
CGATGACGACAAGtaaATAACTTCGTATACTATAGCCTATAGCAATTTATatggcTACCCATACGAT
GTTCCAGATTACGCTtaATAACTTCGTATATACTATACTATAGCAATTTATtaGAACAAAAACTCAT
CTCAGAAGAGGATCTGtaa

| 2) AAV shuttle vectors | | |
|---|---|---|
| Name | Description / Insert | Note |
| pFBAAV-CMV-JT15 | “empty” 5’ vector with a CMV promoter, an MCS, an SD site, and a loxJT15 site | |
| pFBAAV-CMV-JT1522 | “empty” 5’ vector with a CMV promoter, an MCS, an SD site, and a loxJT15:2272 site | |
| pAAV-CBh-JT1522 | “empty” 5’ vector with a CBh promoter, an MCS, an SD site, and a lox15:2272 site | |
| pAAV-CMV-JT15-CRE | “empty” 5’ vector with a CMV promoter, an MCS, an SD site, a loxJT15 site, T2A-CRE, and a BGH polyA signal | |
| pAAV-CMV-15-IRES-CRE | “empty” 5’ vector with a CMV promoter, an MCS, an SD site, a loxJT15 site, an internal ribosome entry site (IRES), CRE, and a BGH polyA signal | |
| pAAV-CBh-JT15-CREv1 | “empty” 5’ vector with a CBh promoter, an MCS, an SD site, a loxJT15 site, T2A-CRE, and a BGH polyA signal | |
| pAAV-CBh-JT15-CREv2 | “empty” 5’ vector with a CBh promoter, an MCS, an SD site (-1 nt shift in the ORF compared to v1), a loxJT15 site, T2A-CRE, and a BGH polyA signal | |
| pAAV-CBh-JT15-CREv3 | “empty” 5’ vector with a CBh promoter, an MCS, an SD site (+1 nt shift in the ORF compared to v1), a loxJT15 site, T2A-CRE, and a BGH polyA signal | |
| pAAV-JTZ17-mid-1522 | “empty” middle vector with a loxJTZ17 site, an SA site, an MCS, an SD site, and a lox15:2272 site | |
| pAAV-JTZ17-mid-15HT1 | “empty” middle vector with a loxJTZ17, an SA site, an MCS, an SD site, and a lox15:HT1site | |
| pAAV-17HT1-mid-1522 | “empty” middle vector with a lox17:HT1 site, an SA site, an MCS, an SD site, and a loxJT15:2272 site | |
| pFBAAV-JTZ17-pA | “empty” 3’ vector with a loxJTZ17 site, an SA site, an MCS, and a polyA signal | |
| pFBAAV-1722-pA | “empty” 3’ vector with a lox17:2272 site, an SA site, an MCS, and a polyA signal | |
| pAAV-EF1$\alpha$-CRE | CRE expression vector with a human EF1$\alpha$ promoter | Figures 2, 3, and 4 |
| pAAV-EF1$\alpha$-DD-CRE | DD-CRE expression vector with a human EF1$\alpha$ promoter | Figures S4 and 7 |
| pAAV-CMV-gp41-IntN | “empty” N-terminal gp41 split intein vector containing a CMV promoter, an MCS, gp41 IntN, and a BGH polyA signal | |
| pAAV-CMV-gp41-IntC | “empty” C-terminal gp41 split intein vector containing a CMV promoter, gp41 IntC, an MCS, and a BGH polyA signal | |
| pFBAAV-CMV-IFT140N-15 | An HA tag and 5’ 1,923 bp of the human IFT140 CDS were inserted into the MCS of pFBAAV-CMV-JT15 | Figures 3 and 4 |
| pAAV-17-BBS1-1522 | 5’ 1,718 bp of the human BBS1 CDS was inserted into the MCS of pAAV-JTZ17-mid-1522 | Figure 3 |
| pFBAAV-1722-LZTFL1-pA | A 3x FLAG tag and the human LZTFL1 CDS (300 bp) were inserted into the MCS of pFBAAV-1722-pA | Figures 3 and 4 |

| | | |
|---|---|---|
| pAAV-17-IFT57-15HT1 | Human IFT57 CDS (1,279 bp) was inserted into the MCS of pAAV-JTZ17-mid-15HT1 | Figure 4 |
| pAAV-17HT1-BBS5-1522 | Human BBS5 CDS (1,022 bp) was inserted into the MCS of pAAV-17HT1-mid-1522 | Figure 4 |
| pAAV-CMV-IFT140N-1522-CRE | IFT140 5’ vector containing a CMV promoter, a HA tag, 5’ 1,923 bp of IFT140 CDS, an SD site, a lox15:2272 site, T2A-CRE, and a BGH polyA signal | Figure 5 |
| pAAV-CBh-IFT140N-1522-CRE | IFT140 5’ vector containing a CBh promoter, a HA tag, 5’ 1,923 bp of IFT140 CDS, an SD site, a lox15:2272 site, T2A-CRE, and a BGH polyA signal | Figure 5 |
| pFBAAV-1722-IFT140C-pA | IFT140 3’ vector containing a lox17:2272 site, an SA site, 3’ 2,466 bp of IFT140 CDS, and a BGH polyA signal | Figure 5 |
| pAAV-CMV-IFT140N-gp41N | An HA tag and 5’ 2,301 bp of the IFT140 CDS (aa1-767) were inserted into the MCS of pAAV-CMV-gp41-IntN | Figure 5 |
| pAAV-CMV-gp41C-IFT140C | 3’ 2,088 bp of the IFT140 CDS (aa768-1462) was inserted into the MCS of pAAV-CMV-gp41-IntC. | Figure 5 |
| pAAV-CMV-secGP41C | The signal peptide sequence of PCDH15 (N-terminal 26 residues) was inserted at the N-terminus of gp41 IntC in pAAV-CMV-gp41-IntC. | |
| pAAV-CMV-PCDH15N-926 | 5’ 2,778 bp of PCDH15 CDS was inserted into the MCS of pAAV-CMV-gp41-IntN | Figure 6 |
| pAAV-CMV-gp41C-PCDH15C-927 | 3’ 3,087 bp of the PCDH15 CDS (with a C-terminal FLAG tag) was inserted into the MCS of pAAV-CMV-secGP41C. | Figure 6 |
| pAAV-CMV-PCDH15N-1035 | 5’ 3,105 bp of PCDH15 CDS was inserted into the MCS of pAAV-CMV-gp41-IntN | Figure 6 |
| pAAV-CMV-gp41C-PCDH15C-1036 | 3’ 2,760 bp of the PCDH15 CDS (with a C-terminal FLAG tag) was inserted into the MCS of pAAV-CMV-secGP41C. | Figure 6 |
| pAAV-CMV-PCDH15N-15-IRES-CRE | 5’ 1,932 bp of PCDH15 CDS was inserted into the MCS of pAAV-CMV-15-IRES-CRE | Figure 6 |
| pFBAAV-17-PCDH15C-pA | 3’ 3,933 bp of PCDH15 CDS (with a C-terminal FLAG tag) was inserted into the MCS of pFBAAV-JTZ17-pA. | Figure 6 |
| pFBAAV-CMV-CEP290N-JT15 | CEP290 5’ vector: 5’ 3,527 bp of CEP290 CDS with an N-terminal FLAG tag was inserted into the MCS of pFBAAV-CMV-JT15 | Figure 7 |
| pFBAAV-17-CEP290C-pA | CEP290 3’ vector: 3’ 3,913 bp of CEP290 CDS was inserted into the MCS of pFBAAV-JTZ17-pA | Figure 7 |
| pAAV-CBh-CEP290-E1-15-CRE | CEP290 5’ vector: a FLAG tag and 5’ 1,065 bp of the human *CEP290* CDS were inserted into the MCS of pAAV-CMV-15-IRES-CRE | Figure 8 |
| pAAV-17-CEP290-E2-1522 | CEP290 middle vector: 2,913 bp from the middle of *CEP290* CDS was inserted into the MCS of pAAV-JTZ17-mid-1522 | Figure 8 |
| pFBAAV-1722-CEP290-E3-pA | CEP290 3’ vector: 3’ 3,462 bp of *CEP290* CDS was inserted into the MCS of pFBAAV-1722-pA | Figure 8 |

| | | |
|---|---|---|
| pAAV-CBh-CDH23-E1-15-CRE | CDH23 5’ vector: an HA tag and 5’ 2,176 bp of the human CDH23 CDS were inserted into the MCS of pAAV-CBh-JT15-CREv2 | Figure 8 |
| pAAV-17-CDH23-E2-1522 | CDH23 middle vector: 4,077 bp from the middle of CDH23 CDS was inserted into the MCS of pAAV-JTZ17-mid-1522 | Figure 8 |
| pFBAAV-1722-CDH23-E3-pA | CDH23 3’ vector: 3’ 3,812 bp of CDH23 CDS was inserted into the MCS of pFBAAV-1722-pA | Figure 8 |

| 3) Other expression plasmids used | | |
|---|---|---|
| Name | Description / Insert | Note |
| Reporter loxP-2272 | Reporter construct to detect recombination of loxP with loxm7, loxHT1, loxHT2, and lox2272 | Figure 2 |
| Reporter loxP-N | Reporter construct to detect recombination of loxP with loxm7, loxHT1, loxHT2, and loxN | Figure 2 |
| Reporter lox2272-N | Reporter construct to detect recombination of lox2272 with loxm7, loxHT1, loxHT2, and loxN | Figure 2 |
| Reporter loxP-HT2 | Reporter construct to detect recombination of loxP with loxm7, loxHT1, and loxHT2 | Figure 2 |
| Reporter lox2272-HT2 | Reporter construct to detect recombination of lox2272 with loxm7, loxHT1, and loxHT2 | Figure 2 |
| pcDNA3-MYC-hBBS1 | Expresses MYC-tagged human BBS1 (WB control; (Seo *et al*, 2009)) | Figure 2 |
| pCS2HA-hLZTFL1 | Expresses HA-tagged human LZTFL1 (WB control; (Seo *et al*, 2011)) | Figure 2 |
| pCS2FLAG-hLZTFL1 | Expresses FLAG-tagged human LZTFL1 (WB control; (Seo *et al.*, 2011)) | Figure 2 |
| pCS2HA-hIFT140 | Expresses HA-tagged human IFT140 | Figure 5 |
| pSS-FS-hCEP290 | Expresses FLAG-tagged human CEP290 | Figure 7 |
| pSS-HA-hCDH23-SF | Expresses HA-tagged human CDH23 | Figure 8 |

**Table S1**. Antibodies used in this study.

| Antibody | Source/Reference | Cat # | Verification |
|---|---|---|---|
| Mouse anti-β-actin (monoclonal; AC-15) | Sigma-Aldrich | A1978 | (Datta *et al*, 2020; Gimona *et al*, 1994) |
| Rabbit CEP290 (polyclonal) | Bethyl Lab | IHC-00365 | (Zhang *et al*, 2014); this study |
| Rabbit CRE (monoclonal; D7L7L) | Cell Signaling | 15036S | this study |
| HRP-linked anti-FLAG (monoclonal; M2) | Sigma-Aldrich | A8592 | N/A |
| HRP-linked anti-HA (monoclonal: 3F10) | Sigma-Aldrich | 12013819001 | N/A |
| Rabbit anti-IFT140 (polyclonal) | ProteinTech | 17460-1-AP | this study |
| Rabbit anti-LZTFL1 (polyclonal) | Seongjin Seo | N/A | (Datta *et al*, 2015; Seo *et al.*, 2011) |
| HRP-linked anti-MYC (monoclonal; 9E10) | Santa Cruz Biotechnology | SC-40 HRP | N/A |
| Sheep anti-PCDH15 (polyclonal) | R&D systems | AF6729 | this study |
| Rabbit anti-V5 (monoclonal: D3H8Q) | Cell Signaling Technology | 13202 | N/A |
| HRP-linked anti-mouse IgG | Cell Signaling Technology | 7076 | |
| HRP-linked anti-rabbit IgG | Cell Signaling Technology | 7074 | |
| HRP-linked anti-sheep IgG | R&D Systems | HAF016 | |
